# Genomic prediction of individual inbreeding levels for the management of genetic diversity in populations with small effective size

**DOI:** 10.1101/2024.05.31.594735

**Authors:** Natalia Soledad Forneris, Mirte Bosse, Mathieu Gautier, Tom Druet

## Abstract

In populations of small effective size (N_e_), such as those in conservation programs, companion animals or livestock species, management of diversity and inbreeding is essential. Homozygosity-by-descent (HBD) segments provide relevant information in that context, as they allow efficient estimation of the inbreeding coefficient, provide locus-specific information and their length is informative about the “age” of inbreeding. Therefore, our objective was to evaluate tools for predicting HBD in future offspring based on parental genotypes, a problem equivalent to identifying segments identical-by-descent (IBD) among the four parental chromosomes. In total, we reviewed and evaluated 16 approaches using simulated and real data with small N_e_. The methods included model-based approaches, mostly hidden Markov models (HMMs), which considered up to 15 IBD configurations among the four parental chromosomes, as well as more computationally efficient rule-based approaches. The accuracy of the methods was then evaluated, including with low-density marker panels, genotyping-by-sequencing data and small groups of individuals, typical features in such populations. Two HMMs performed consistently well, while two rule-based approaches proved efficient for genome-wide predictions. The model-based approaches were particularly efficient when information was reduced (low marker density, locus-specific estimation). Methods using phased data proved to be more efficient, while some approaches relying on unphased genotype data proved to be sensitive to the allele frequencies used. In some settings, pedigree information was competitive in predicting recent inbreeding levels. Finally, we showed that our evaluation is also informative about the accuracy of the methods for estimating relatedness and identifying IBD segments between pairs of individuals.

## Introduction

In conservation genetics, companion animals and livestock species, effective population sizes (N_e_) are often small and optimal management strategies are implemented to maintain genetic diversity and limit levels of inbreeding, often relying on reproducer selection and mating advice. The expected level of inbreeding of future offspring is an important criterion for such mating advice, as it allows the risk of inbreeding depression (ID) or genetic defects to be reduced.

The inbreeding coefficient (*F*) of an individual has been defined as the correlation between the uniting gametes (Wright, 1922) and the probability that the two alleles present at a locus are identical-by-descent (IBD), i.e. inherited twice from a common ancestor (Malécot, 1948). In that case, the neighboring loci will also be IBD because a whole segment has been inherited IBD from the founder. In the absence of mutations, these segments are homozygous and are therefore called homozygous-by-descent (HBD) (Schäffer, 1999). The length of these HBD segments depends on the number of generations *G* to the common ancestor, with more generations providing more opportunities for the recombination process to cut the transmitted segment. With genotyping data, these HBD segments will appear as long stretches of homozygous genotypes called runs-of-homozygosity (ROH), which are often used as a proxy for HBD and to estimate *F* (Broman and Weber, 1999; McQuillan et al., 2008). Model-based approaches have also been implemented to describe individual genomes as a mosaic of HBD and non-HBD segments (Leutenegger et al., 2003), allowing identification of HBD segments, calculation of HBD probabilities at each marker position, and estimation of realized inbreeding levels. Such model-based approaches have been shown to be more efficient than rule-based methods when information is degraded or marker density is low (Druet and Gautier, 2017; Lavanchy and Goudet, 2023).

ROH-based (*F*_ROH_) and HBD-based (*F*_HBD_) estimators of *F* present several advantages over other marker-based estimators. Indeed, studies from Nietlisbach et al. (2019) and Caballero et al. (2021) showed that *F*_ROH_ performs well in populations with low N_e_. In agreement, Alemu et al. (2021) concluded that HBD-based methods are efficient in livestock species where deleterious alleles can reach high frequencies (e.g., Keller and Waller, 2002; Bosse et al., 2019). In addition, these methods provide locus-specific estimates, which can be used to manage recessive alleles that cause genetic defects or have a large contribution to ID. They are also informative about the age of HBD segments (Kirin et al., 2010; Pemberton et al., 2012), thus allowing to estimate the recent inbreeding that is expected to be deleterious (Hinrichs et al., 2007; Szpiech et al., 2013; Stoffel et al., 2021; Naji et al., 2024) and therefore more relevant for management strategies. They are also more robust to the used allele frequencies (AF), that might introduce biases (Keller et al., 2011; Caballero et al., 2022; Naji et al., 2024). Finally, these estimators are more interpretable as they range between 0 and 1 as the pedigree-based estimators, and allow to define a base population comparable to the pedigree (Solé et al., 2017). In summary, the use of measures based on HBD segments offers several advantages for managing diversity and inbreeding in populations with small N_e_, as in conservation genetics, some wildlife species, companion animals and livestock populations. Actually, the benefits of using segment-based measures to maintain diversity and fitness have been demonstrated in similar populations (de Cara et al., 2013; Bosse et al., 2015; Gómez-Romano et al., 2016), for example by minimizing the occurrence of long HBD segments in offpsring (Bosse et al., 2015). Similarly, Meuwissen et al. (2020) concluded that the use of IBD-based approaches performed well in optimal contribution selection schemes.

In the present study, our objective was to evaluate different methods for predicting inbreeding levels in future offspring based on genotypes from their parents, as these are an important component in mating plans implemented to manage diversity and inbreeding in populations with small N_e_. More specifically, our aim was to predict future HBD levels, as these metrics have been proven to be efficient and offer several advantages (see above). Therefore, we reviewed methods for identifying IBD segments between parental chromosomes and predicting HBD levels in offspring, and selected state-of-the-art model– and rule-based approaches to perform a comprehensive evaluation study. Importantly, this evaluation was performed on real data sets from a large sequenced cattle pedigree and a population of Mexican wolves, similar to those typically found in the field of conservation genetics, animal breeding or molecular ecology. It is indeed essential to measure their accuracy in such settings because the genomic structure of the population has been shown to affect their performance (e.g. Caballero et al., 2021), while most of the methods have only been evaluated on human datasets characterized by large N_e_. Simulated data with similar population structures were also generated to consolidate our results. Accuracy was also assessed at lower marker density or with genotyping-by-sequencing (GBS) data, genotyping strategies that are frequently used in these populations. We start by a description of the different evaluated methods.

## Material and methods

### Overview of approaches to predict homozygosity-by-descent in offspring

Our objective is to predict the level of inbreeding of an individual based on the genotypes of its parents (Figure 1A). More specifically, the aim is to predict the proportion of HBD segments and to estimate the probability that a locus will be HBD. The expected inbreeding coefficient of an individual is equal to the kinship coefficient between its two parents (Malécot, 1967). Similarly, the prediction of HBD can be obtained from the probability that the haplotypes transmitted by the parents are IBD. Each parent can transmit two haplotypes, so there are four possible combinations of pairs of parental haplotypes at a locus, and the HBD probability can be computed from the IBD probabilities of these four combinations (Figure 1). Note that due to recombination, the transmitted haplotypes might differ from the parental haplotypes. Nevertheless, the expected HBD level can still be obtained as the average IBD of the four parental combinations (recombination rates and interference levels do not affect the expected inbreeding levels but their variance).

**Figure 1.**
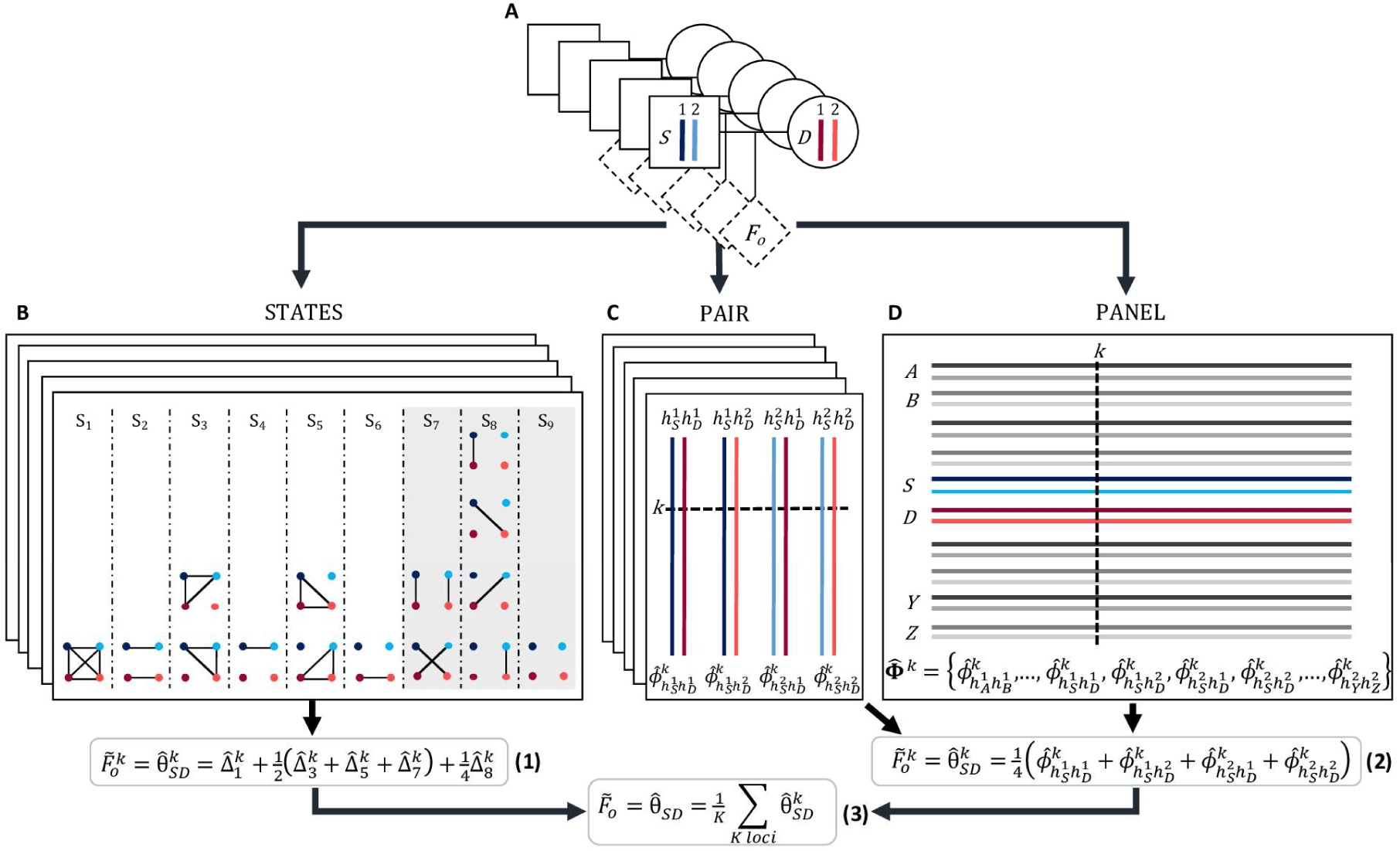
Approaches used to predict the homozygous-by-descent (HBD) level of an individual (*F*_O_) based on the genotypes of its parents. A) Predictions were made in several trios, each including a sire (S) and a dam (D), having each a paternal and maternal haplotype (labelled 1 and 2 respectively), shown in blue for the sire and red for the dam, and the future offspring (dashed diamond). B) STATES approach: for each pair of parents, the four parental haplotypes can take fifteen possible identity-by-descent (IBD) configurations at a locus (Jacquard, 1974). The four parental haplotypes are represented by dots using the same colors as in A) while solid lines indicate IBD between linked haplotypes. The 15 IBD states can be grouped into 9 condensed IBD states {*S*_1_, *S*_2_, …, *S*_9_} if the parental origins of the haplotypes within the two individuals are considered unknown. The number of states are reduced to three if the parents are assumed to be non-inbred (grey background). The STATES approaches model the observed genotypes or haplotypes conditional on these configurations. At locus *k*, the estimated probabilities of the 9 IBD modes 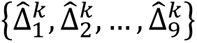 can be used to estimate the locus specific coancestry between the parents 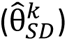, which corresponds to the predicted locus-specific HBD level in the offspring 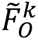 (Eq. 1). C) PAIR approach: IBD is modelled sequentially for each of the four possible combinations of parental haplotypes 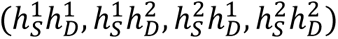, where 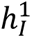 and 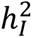 denote the paternal and maternal haplotypes of individual *I*, respectively. This analysis estimates the IBD probability 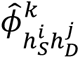 between two haplotypes 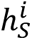 and 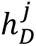 at locus *k* for the four possible pairs. D) PANEL approach: a large panel of haplotypes from a large number of individuals are analyzed jointly to detect IBD segments. At locus *k*, a vector 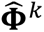 containing the IBD probabilities for each pair of haplotypes is estimated (note that with rule-based approaches, IBD probabilities are either 0 or 1). In the PAIR and PANEL approaches, the locus specific coancestry between the parents is obtained as the average over the four possible pairs of parental haplotypes (Eq. 2). Finally, the genome-wide HBD levels 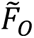 can the predicted as the average locus-specific values at the *K* loci (Eq. 3).

The methods we will evaluate are therefore based on the modeling of the IBD relationships between the four chromosomes of two individuals (here, the four parental homologues) that can take on fifteen different configurations (Figure 1B), called the identity states (Jacquard, 1974). If the parental origins of the haplotypes within the two individuals are unknown, symmetric configurations are equivalent and the 15 identity-states can be grouped into 9 condensed identity-states (Jacquard, 1974). The configurations can be further reduced to three states if the inbreeding of the individuals is ignored. In this case, the three states simply correspond to the sharing of 0, 1 and 2 IBD haplotypes between the two individuals. Accordingly, several methods model the observed genotypes or haplotypes conditionally on these possible configurations (referred to as 15, 9 or 3-STATES approaches). Another approach is to model chromosomes individually, ignoring which chromosomes belong to the same individual. In this case, methods can either model the IBD relationship between each possible pair of haplotypes sequentially (PAIR approach – Figure 1C), leading to the analysis of four pairs when working with two individuals, or simultaneously model all haplotypes from all individuals in the analyzed sample and efficiently identify IBD sharing in this panel of haplotypes (PANEL approach – Figure 1D).

Prediction methods can be further classified according to whether they use genotypes or haplotypes. The use of genotypes is only possible in the 9 or 3-STATES approaches by modeling the genotype probabilities conditionally on the underlying state. Haplotypes require phasing of the data, a procedure that provides additional information, but that can also introduce errors.

Finally, we can also distinguish between rule-based and model-based approaches. Rule-based approaches use a set of rules to determine whether two individuals share IBD haplotypes based on their genotypes or whether two haplotypes are IBD. These rules are typically based on the number of identical-by-state (IBS) alleles or genotypes, the length of the segments compared, or the number of mismatches. Model-based approaches compute the probability of different configurations based on the likelihood of observing the data conditional on the configuration. The possible configurations could, for example, correspond to the 9 identity states in a 9-STATES model, or to IBD versus non-IBD for a PAIR approach. Estimation of these likelihoods may include parameters such as the AF, the probability of genotyping error, the genetic distance between successive markers, etc. Model-based approaches can also handle genotype probabilities to account for genotype uncertainty, e.g. with low-fold sequencing data.

### Description of the evaluated IBD estimation methods

#### STATES approaches

IBD_Haplo (Thompson, 2008; 2009), GIBDLD (Han and Abney, 2011; 2013) and LocalNgsRelate (Severson et al., 2022) model the IBD process along the four parental chromosomes based on continuous time Markov chains that can be implemented as HMM. In these three methods, the emission probabilities are based on AF and on genotyping error probabilities or genotyping uncertainty, and the transition probabilities are a function of the genetic distances. Through local decoding, they estimate the posterior state probabilities, P(X_i_=*k*), at each locus *i*, where X_i_ is the unknown state at locus *i* and *k* is one of the modeled states. With genotyping data, these hidden states correspond to the 9 condensed identity states with IBD_Haplo and GIBDLD, and to the sharing of 0, 1 or 2 IBD with LocalNgsRelate (3-STATES model). With haplotype data, IBD_Haplo fits a model with 15 hidden states corresponding to the 15 identity-states. At each locus, the offspring HBD can be predicted from the locus-specific state probabilities P(X_i_=*k*) using the rules shown in Figure 1. The three methods define the transition matrices differently. For example, in IBD_Haplo they are based on two parameters related to N_e_ and the number of generations to the common ancestors. The model assumes that only one additional IBD relationship can be gained or lost between two successive markers. These parameters are not estimated and are identical for all pairs of individuals. Conversely, GIBDLD and LocalNgsRelate estimate the parameters for each pair of individuals. These parameters include the proportions of the genome associated with the different condensed identity-states, which also provide a genome-wide estimate of relatedness. They also present differences in the emission probabilities. GIBDLD accounts for LD by allowing the emission probabilities at a given locus to depend on the genotypes at a defined number of preceding loci. LocalNgsRelate is designed to work with low depth sequencing data, a feature made possible by using genotype likelihoods to define emission probabilities. Like LocalNgsRelate, TRUFFLE (Dimitromanolakis *et al*., 2019) fits a 3-STATES model but uses a rule-based approach.

#### PAIR approaches

When modeling IBD for pairs of haplotypes, it is possible to use methods designed to identify ROH or HBD segments within individual genomes. Although this is somewhat artificial, since the two haplotypes don’t really belong to the same individual, it has the advantage that they behave like the HBD measures we want to predict in the offspring. This strategy has been used, for example, with ROH in several studies (Pryce et al., 2012; Bosse et al., 2015; de Cara et al., 2013). The ZooRoH model (Druet and Gautier, 2022) is an alternative to rule-based approaches to identify ROH. It is an HMM that describes the genome as a mosaic of HBD and non-HBD segments. A specific feature of ZooRoH is that it fits several HBD classes, each class *c* having its own rate parameter *R*_c_ that defines the expected length of HBD segments (equal to 1/R_c_ Morgans). Each class is therefore associated with a different set of ancestors present in different past generations. This makes it possible to estimate the level of inbreeding with respect to different base populations (Solé et al., 2017). Note that when a single HBD class is fitted, we refer to a ZooRoH-1R model. This model is identical to that of Leutenegger et al. (2003) and has only two parameters, a rate and a mixing coefficient, both estimated for each individual.

#### PANEL approaches

GERMLINE (Gusev et al., 2009), hap-IBD (Zhou et al., 2020) and phasedibd (Freyman et al., 2021) belong to the PANEL approach and use rule-based methods to find long segments shared IBS between two haplotypes. These three methods avoid comparing each pair of haplotypes sequentially to improve their computational efficiency. For this purpose, GERMLINE is based on hash tables and libraries of haplotypes, while hap-IBD uses the Positional Burrows-Wheeler Transform (PBWT; Durbin, 2014) and phasedibd relies on an extension of the PBWT called the Templated PBWT, which allows to mask errors and expand haplotype matches. Although they share a common principle, these methods differ in their implementation and their handling of genotyping and haplotyping errors. For example, hap-IBD and phasedibd can account for genotyping errors and even correct for some so-called switch errors resulting from the phasing process. Finally, Refined-IBD (Browning and Browning, 2013) first relies on GERMLINE to identify long segments shared IBS, and then uses a LOD score to determine whether the haplotypes are IBD or not.

#### Pedigree-based and genomic relationship matrices

The genome-wide inbreeding coefficient can also be predicted as half the relationship between the parents obtained using either genealogical information or genotyping data, SNP-by-SNP, and the rules described in VanRaden (2008). Depending on the selected rules, using the elements from the genomic relationship matrix (GRM) is equivalent to predicting F_GRM_ (VanRaden, 2008) or the correlation between the uniting gametes, F_UNI_ (Li and Horvitz, 1953; Yang et al., 2010).

### Evaluation design

The objective of the present study is to evaluate the accuracy of predicting HBD in future offspring based on parental genotypes in populations with small N_e_. For this evaluation we used both simulated and real data sets. With simulations, the true HBD levels are known, whereas with real data, the methods are evaluated in more realistic data structures and conditions. Therefore, we used a design that can be applied to both datasets, taking advantage of the genotyped trios available for the real data. In this design, we use the parental genotypes to perform the predictions and the offspring genotypes to evaluate their accuracy (Figure S1). The genotypes of the offspring provide the possibility to estimate its realized inbreeding. In particular, with sequence data, genotypes are available for all variants, allowing accurate estimation of whole-genome heterozygosity (Kardos et al., 2016; Alemu et al., 2021).

The accuracy was then assessed using the correlations between the predicted and reference genome-wide levels of HBD (defined as the average locus-specific HBD levels). For locus-specific predictions, the correlations between predicted and reference locus-specific levels of HBD were computed, and ROC curves associated with these predictions were derived.

### Software and parameters used for prediction of homozygosity-by-descent

For evaluation, offspring genotyping data were first removed from the analysis. Parental genotypes were then phased using Beagle 5.4 (Browning et al., 2021) and default parameters. The software used to predict global and locus-specific HBD levels in offspring based on parental genotypes or haplotypes are listed in Table 1 and Table S1. For IBD_Haplo, GIBDLD, LocalNgsRelate and TRUFFLE, we essentially used default parameters (see Table S1). IBD_Haplo (Thompson, 2008) was run either with genotypes (IBD_Haplo9c), either with haplotypes (IBD_Haplo15c) and the transition matrix described in Brown et al. (2011). For RZooRoH (Bertrand et al., 2019), we used a ‘layer’ model (Druet and Gautier, 2022) with 6 HBD classes with predefined rates R_c_ = {5, 25, 125, …, 15,625}. We then estimated IBD between each pair of parental haplotypes using posterior probabilities from HBD classes with R_c_ ≤ 25 (recent IBD – ZooRoH-25) or from HBD classes with R_c_ ≤ 125 (total IBD or ZooRoH-125). The last HBD classes were not included as they are less reliable (only a few SNPs per segment) and more dependent on AF like maximum likelihood estimators (Alemu et al., 2021). In addition, we also ran a model estimating the rate of a single HBD class (ZooRoH-1R). ROH were identified using PLINK (Purcell et al., 2007) with 40 SNP windows and minimum length of 2 Mb. We also used default parameters for Refined-IBD, GERMLINE and hap-IBD, while the minimum segment length was set to 2 Mb for phasedibd (Table S1).

**Table 1.**
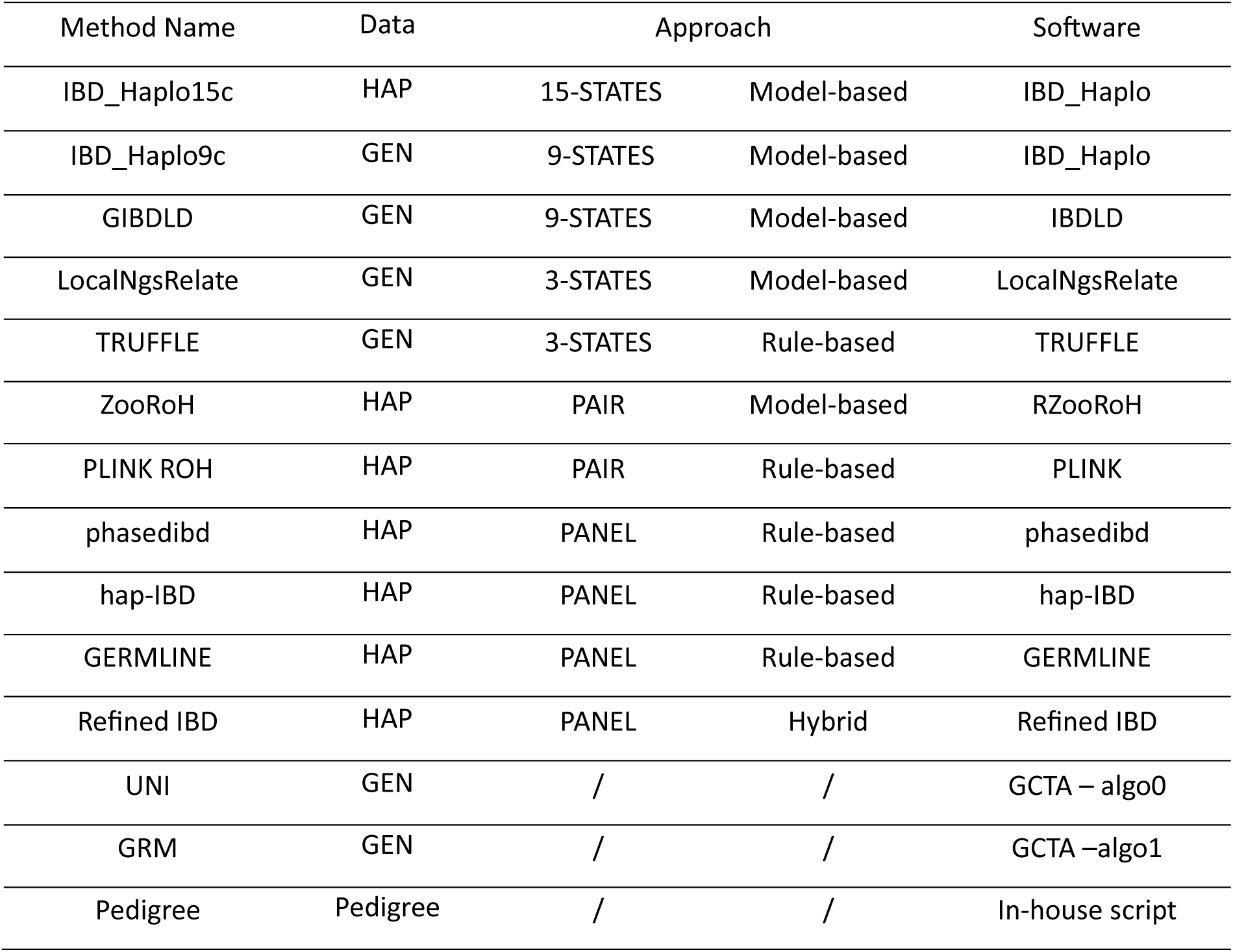
Main properties of the methods used to predict levels of homozygosity-by-descent in a future offspring of a genotyped pair of parents. The table also indicates whether the methods use genotype (GEN) or haplotype (HAP) data and the names of the corresponding software. The STATES, PAIR and PANEL approaches are described in the main text and in Figure 1.

By default, methods use AF estimated from the sample, although ideally the AF of the base population should be used. Therefore, we also tested methods that accept external AF as input (IBD_Haplo, ZooRoH, LocNgsRelate, GRM and UNI) with founder AF estimated using the gene content approach of Gengler et al. (2007). SNPs with a founder MAF less than 0.005 were discarded.

### Data

#### Simulation study

We simulated two populations with small N_e_ in the most recent generations. These demographic histories, characterized by successive reductions in N_e_, were chosen to obtain populations with similar levels of inbreeding and genomic structure to the two real datasets (see below). The scenarios resulted in moderate and high levels of inbreeding, termed MODF and HIGHF, corresponding to the levels observed in a typical livestock population (dairy cattle) and an endangered wild population (the Mexican wolf).

Simulations were performed with SLiM 4.0.1 (Haller and Messer, 2023) and msprime 1.2.0 (Baumdicker et al., 2022) using forward-time simulation and a “recapitation” technique (Haller et al., 2018). For both populations, individual genomes consisted of 25 chromosome pairs of 100 cM each. Recombination and mutation rates were set to 10^-8^ per bp. The initial population consisted of 10,000 diploid individuals, mating was assumed to be random and the sex ratio was set to 1. Subsequent evolution of demographic parameters is described in Figure 2. In total, 100 and 20 offspring were simulated in the last generation in the MODF and HIGHF scenarios respectively. The simulated data consisted of the trios formed by these offspring and their parents. In each replicate, we randomly selected a subset of 5,000 or 25,000 evenly spaced bi-allelic markers with a MAF ≥ 0.01 (corresponding to low and medium-density arrays with 1 SNP / cM and 10 SNPs / cM). In addition, we kept track of the pedigree for the last 15 generations. Each scenario was repeated 100 times.

**Figure 2.**
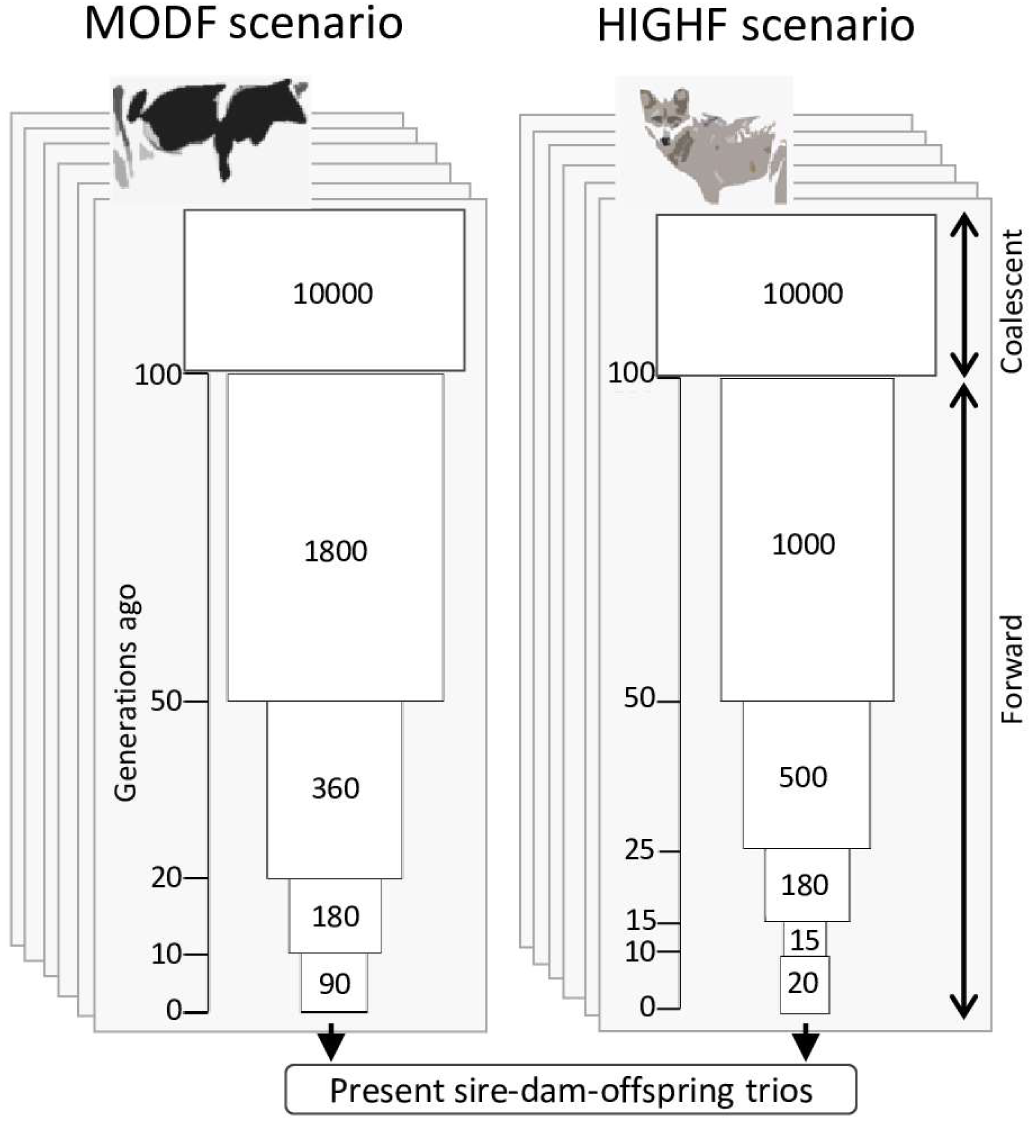
Simulated demographic scenarios. We simulated a moderately inbred (MODF) population corresponding to a typical livestock population (e.g. dairy cattle), and a highly inbred (HIGHF) population corresponding to an endangered wild population under conservation (similar to the Mexican Wolf). In both scenarios, we run a forward-in-time simulation with an initial population of N_e_ = 10,000 and equal sex ratio. In the MODF and HIGHF scenario, the proportion of males for the subsequent generations was set to 0.1 and 0.5, respectively. The figure represents the evolution of N_e_ across generations.

We used the tree sequence recording feature to keep track of the true local ancestry at each marker position for each individual. At each position, HBD was then declared if the time to the most recent common ancestor (TMRCA) was less than the age of the defined base population expressed in generations. We used three different base populations to define the reference levels of inbreeding, corresponding respectively to recent inbreeding only (young base population set to 15 generations – F_YOUNG_), to 50 generations of inbreeding (intermediate base population – F_MID_), and to a large number of generations including ancient inbreeding (ancient base population set to 500 generations – F_TOT_).

#### Dutch Holstein data set

The first real data set represents a population typical of livestock species, with a small N_e_ and under intense selection. We used whole-genome sequence data (8,417,679 SNPs) available for 264 Dutch Holstein individuals from the DAMONA cattle pedigree previously used by Oget-Ebrad et al. (2022). This pedigree can be divided into 127 parents, corresponding to 84 different pairs, and their 98 offspring. Offspring belonging to a sequenced trio that were also parents in another trio, were excluded from the list of sequenced parents. The pedigree available for these individuals consisted in 12,238 individuals.

The reference inbreeding measures were computed using ZooRoH and the whole-genome sequence data because inbreeding estimation with ZooRoH proved highly accurate on the simulated data (Table S2). In addition, the model has been proven efficient and robust to genotyping errors in previous studies. We ran a ‘layer’ model (Druet and Gautier, 2022) with 6 HBD classes with fixed rates R_c_ = {5, 25, 125, 625, 3125, 15,625} on the sequence data from the offspring only. Recent inbreeding F_YOUNG_ was defined using only posterior HBD probabilities of classes with R_c_ ≤ 25, while F_MID_ and F_TOT_ were obtained using HBD classes with R_c_ ≤ 125 and R_c_ ≤ 3125, respectively. These correspond approximately to base populations set at 12.5, 62.5 and 1500 generations in the past.

#### Mexican Wolf data set

We also evaluated the accuracy of the methods in a population with higher levels of inbreeding and under a conservation program. It consisted of 13 trios from the endangered Mexican Wolf (MW, *Canis lupus baileyi*) population genotyped using the Illumina CanineHD BeadChip array (Illumina, Inc., San Diego, CA). The MW population was reintroduced into the wild in the 1990s. It is derived from three unrelated captive lineages, called McBride (MB), Aragón (AR) and Ghost Ranch (GR), each descended from 2 or 3 founders. The trios were extracted from a data set (Fitak *et al.,* 2018) including 88 MW genotyped for 118,287 SNPs. One individual was excluded for having more than 10% Mendelian conflicts. In addition to the original filtering, we kept only markers with call rate ≥ 0.90, MAF ≥ 0.01, without Mendelian conflicts and in Hardy-Weinberg equilibrium (p ≥ 0.05). The final set included 33 MB, 2 AR, 6 GR and 46 crossbred (i.e. crosses between two lineages) individuals genotyped for 54,037 SNPs. Of these crossbred individuals, 13 individuals and their genotyped parents formed the trios used in our validation experiment. The reference inbreeding measures were computed using all the selected SNPs and the same approach as for the DAMONA data set.

### Impact of marker density and genotyping method

The methods were evaluated with different marker panels, such as low and medium genotyping densities (LD and MD arrays). For the simulated data sets, these corresponded to densities of 1 and 10 SNPs per Mb. For the DAMONA data set, we selected markers in common with the Illumina BovineLD and BovineSNP50 commercial arrays. This resulted in a selection of 5,388 and 29,375 SNPs respectively (approximately 2 and 10 SNPs per Mb). In addition, the DAMONA sequence data allowed us to mimic genotyping-by-sequencing (GBS) data. To do this, we performed an *in silico* digestion of the bovine reference genome using the *Pstl* restriction enzyme using the GBSX package (Herten *et al*., 2015). We selected 72,828 fragments out of 1,503,470 fragments with a length between 200 and 300 bp. The distance between fragments is shown in Figure S2. A total of 60,842 SNPs from the WGS data were located in the selected fragments. To account for allelic dropout (Gautier et al., 2013), when a SNP was located near the restriction site (± 3bp), heterozygous individuals were considered homozygous in the associated fragment for the haplotype carrying the reference alleles (the other haplotype was not amplified). If individuals were homozygous for the alternate alleles of the SNP in the restriction site, all genotypes in the associated fragment were set to missing. We then filtered out SNPs surrounding the restriction site (± 3bp), those with more than 5% missing genotypes or MAF < 0.01. This resulted in a GBS panel of 56,098 SNPs (GBS-50K). To further reduce the number of markers, we first selected fragments with a length ranging between 250 and 300 bp, resulting in a panel of 31,339 SNPs (GBS-30K). A smaller panel of 15,493 SNPs (GBS-15K) was obtained by randomly sampling half of these fragments. Finally, we used only one marker density for the MW (54,037 SNPs, corresponding to about 20-25 SNPs per Mb). In addition to their variable marker densities, the different marker panels differ in the distribution of their AF and of their marker spacing (Figure S3). These elements could also influence the properties of the prediction methods.

## Results

The simulated data closely matched the inbreeding levels and partitioning of HBD observed in the corresponding real data (Figure S4). Comparisons also illustrate that dairy cattle and Mexican wolf populations, and the corresponding MODF and HIGH simulations, provide complementary scenarios.

## Accuracy of predicted genome-wide HBD levels

We first evaluated the predictions obtained using MD genotyping arrays. The correlations between predicted and reference levels of HBD for the 16 methods compared and the four scenarios are shown in Figure 3 and Figure S5 (significance levels of pairwise comparisons are available in Table S3). The reference levels of HBD were defined with respect to different base populations, F_YOUNG_, F_MID_ and F_TOT_, corresponding to approximately 15, 50 and 500 generations. In the simulations, the correlations were high, ranging from 0.78 to 0.86 and from 0.74 to 0.83 in the MODF and HIGHF scenarios, respectively. The ranking of the methods was comparable in the two simulation scenarios and for the different base populations. Several methods, including IBD_Haplo15c, ZooRoH, phasedibd, PLINK ROH and hap-IBD, were consistently among the best methods. Using unphased data and a 9-STATES model with IBD_Haplo (IBD_Haplo9c) also performed well, but was generally less accurate than using phased data and a 15-STATES model. For ZooRoH, only minor differences were observed between the three models tested. Among the three best rule-based approaches, the correlations obtained with hap-IBD were generally slightly lower than those obtained with phasedibd or PLINK-ROH. On the other hand, GIBDLD and LocalNgsRelate achieved lower correlations in the model-based group, while this was the case for GERMLINE and TRUFFLE in the rule-based group. Refined-IBD had one of the lowest correlations in most scenarios. As with GIBDLD and LocalNgsRelate, the relative performance of the SNP-by-SNP approaches was rather variable. Finally, the vast majority of genotype-based predictors outperformed the pedigree-based predictions.

**Figure 3.**
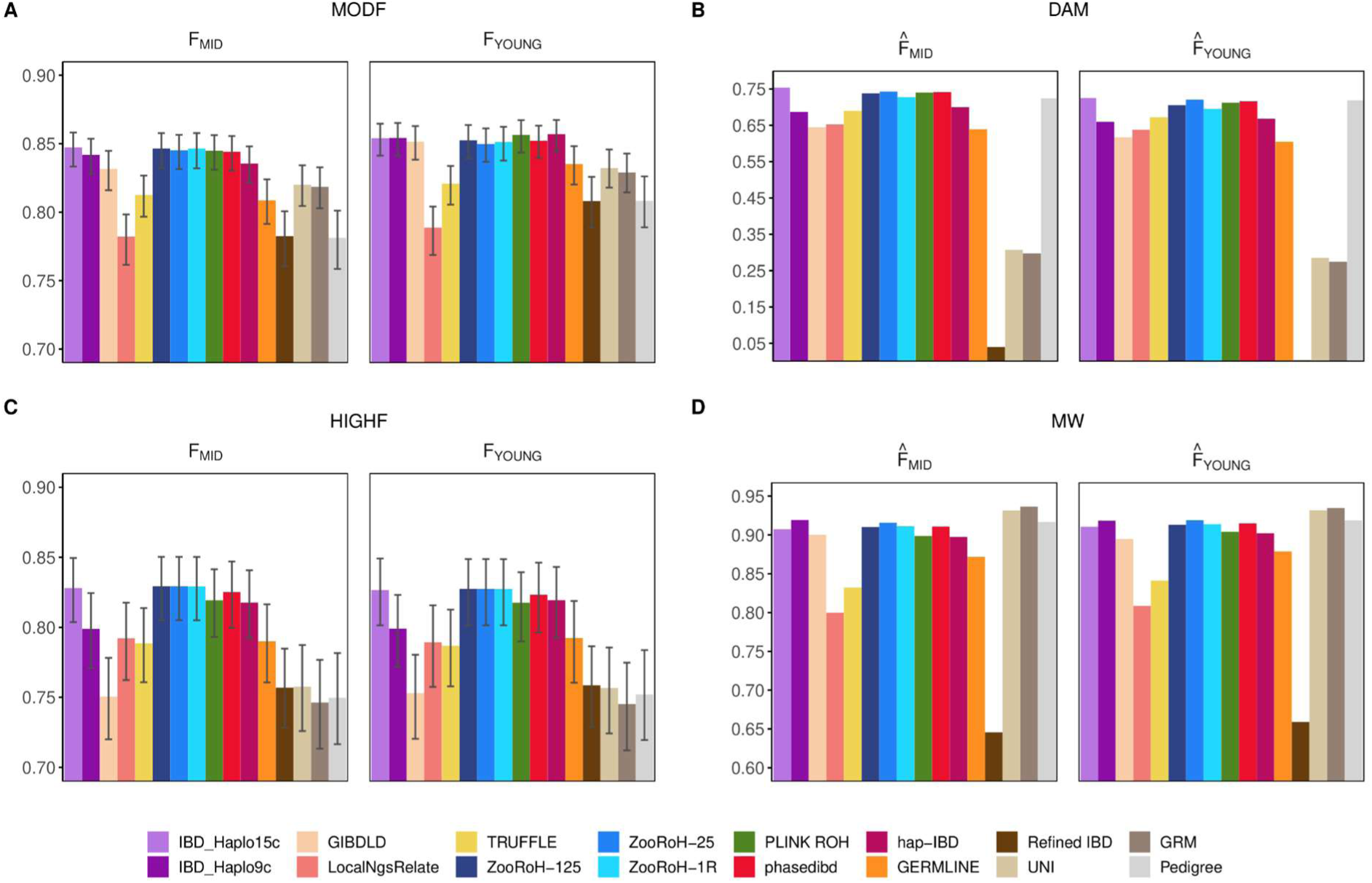
Correlations between predicted and reference genome-wide levels of HBD for the 16 methods compared and the four scenarios, using a medium density array. Methods and their abbreviation are described in Table 1. A) moderately inbred simulated population (MODF); B) DAMONA cattle data set (DAM); C) highly inbred simulated population (HIGHF); D) Mexican Wolf data set (MW). The reference levels were defined for either a recent or an intermediate base population (approximately 15 or 50 generations ago, respectively). Mean and 99% confidence intervals are shown for the simulated scenarios.

Compared to the simulations, the correlations between predicted and reference HBD levels were lower in the DAMONA data set, which contains about 100 trios for evaluation, and higher in the MW population, where we have only 13 trios. Regarding the relative performance of the evaluated methods, some trends observed in these two real data sets were similar to those highlighted with simulated data. IBD_Haplo15c, ZooRoH, PLINK ROH and phasedibd performed well, closely followed by Hap-IBD. Conversely, Refined-IBD, TRUFFLE and LocalNgsRelate had poor relative performance in at least one scenario. GERMLINE and GIBDLD were closer to the best methods, but still achieved systematically lower correlations than, for example, IBD_Haplo15c or phasedibd. As in the simulated data set, IBD_Haplo15c performed better than the 9-STATES model in the cattle data, while the performances of different ZooRoH models were close. The SNP-by-SNP approaches still had variable performances, achieving the highest correlations in the MW population but almost the worst prediction in the DAMONA data set. Interestingly, the correlations obtained with the pedigree-based approach were quite high, among the best methods, especially for recent HBD levels.

## Accuracy of predicted locus-specific HBD levels

We then evaluated the methods in terms of the accuracy of locus-specific correlations (Figure 4, Figures S6-7). The SNP-by-SNP-based and pedigree-based methods were not included in these comparisons because they only provide genome-wide predictions. As expected, prediction accuracies decreased compared to genome-wide predictions averaged over many more loci. Overall, the best methods were almost the same as for genome-wide predictions. However, the best model-based approaches (IBD_Haplo15c and ZooRoH models) now outperformed the best rule-based methods (phasedibd and PLINK ROH). Although hap-IBD was still competitive in some scenarios, it often achieved less accurate predictions than phasedibd and PLINK ROH. With regard to the model-based approaches, the disadvantage of using unphased data (IBD_Haplo9c versus IBD_Haplo15c) was more pronounced, while the three ZooRoH models still had similar performances. Interestingly, the prediction of recent HBD (ZooRoH-25) was more accurate at predicting HBD levels defined with respect to a recent base population (F_YOUNG_ or F_MID_), which was not clearly observed with genome-wide predictions. As before, the other two model-based approaches, GIBDLD and LocalNgsRelate, continued to show variable performance. The other evaluated methods achieved lower accuracies, for instance systematically lower than hap-IBD. These observations were consistent across all evaluated scenarios with both simulated and real data sets, with a few rare exceptions.

**Figure 4.**
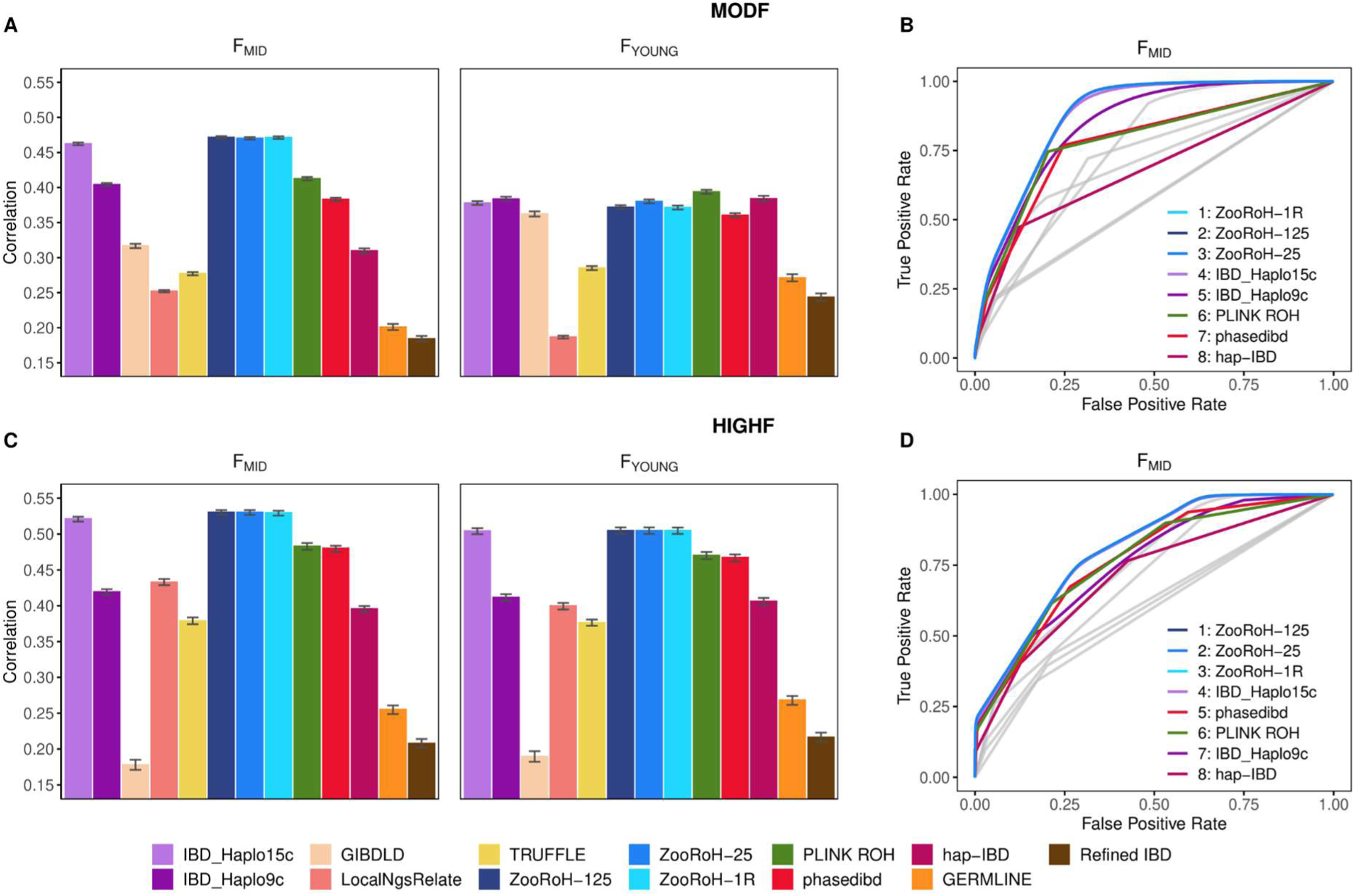
Locus-specific accuracy of 13 HBD prediction methods in the simulated data sets using a medium density array. Methods and their abbreviation are described in Table 1. Accuracy was assessed in the MODF (A-B) and HIGHF (C-D) scenario using correlations (mean and 99% confidence intervals) between predicted and reference locus-specific HBD levels (A & C). The reference levels are the true HBD status at every marker position and defined using either a recent (15 generations ago – F_YOUNG_) or an intermediate (50 generations – F_MID_) base population. The Receiver operating characteristic (ROC) curve for each method were also computed (B & D), with colored curves for the eight best methods. Their relative ranking in terms of AUC is also specified.

When the locus-specific accuracy was assessed using ROC curves (Figure 4, Figure S8) and the associated AUC (Figure S9-10), these trends were confirmed. The advantages of the best model-based approaches were even clearer. ZooRoH and IBD_Haplo15c were indeed associated with the highest AUC across all scenarios. Most often, ZooRoH models were the best, especially when the base population was more ancient. For more recent reference HBD levels, predictions using ZooRoH-25 (recent HBD only) were better than those using ZooRoH-125.

## Accuracy of HBD predictions with low-density marker panels and genotyping-by-sequencing data

To reduce genotyping costs, low-marker density panels or genotyping-by-sequencing (GBS) data are commonly used in genomic studies of wild populations, conservation genetics or livestock species. To evaluate HBD predictions in such data, we relied mainly on the DAMONA data, for which low-marker density panels are available (Boichard et al., 2012) and for which we could use the reference genome to generate *in-silico* GBS panels. In addition, we also defined a LD density panel in the simulations.

For genome-wide HBD predictions using an LD array, accuracy was lower compared to values obtained using an MD genotyping array (Figure 5, Figure S11-12). However, some methods were more robust to this change and their loss of accuracy was limited. This was the case for the best model-based approaches, IBD_Haplo15c and ZooRoH. In the DAMONA data set, some rule-based methods performed even better, including phasedibd, hap-IBD, GERMLINE (with modified parameters) and Refined-IBD. However, this was only confirmed for hap-IBD and GERMLINE in the simulated data. With our parameter setting, no ROH were identified with LD panels (even when reducing the number of SNPs per window and per ROH to 25 and setting the number of heterozygous SNPs to 0). The overall ranking was not much affected, with IBD_Haplo and ZooRoH still among the best methods in all scenarios, while among the rule-based methods, hap-IBD was more reliable. Interestingly, the pedigree-based predictions ranked better and were closer to the best methods at this marker density. Overall, the best methods were still efficient when using a LD marker panel, resulting in correlations close to those obtained using a MD marker panel, especially for recent inbreeding.

**Figure 5.**
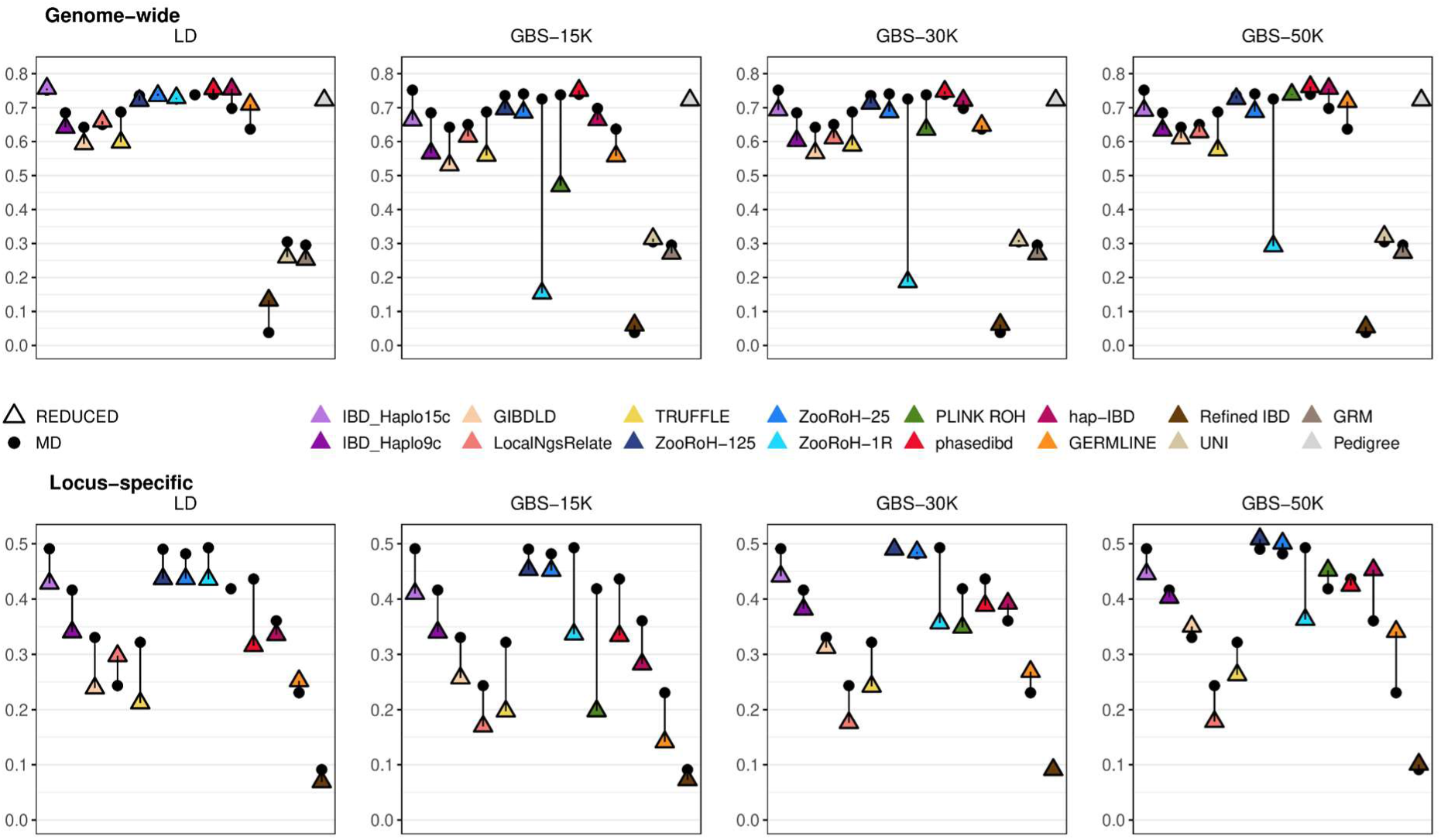
Correlations between predicted and reference genome-wide or locus-specific HBD levels for the 16 evaluated methods in the DAMONA cattle data set using reduced genotyping arrays. Methods and their abbreviation are described in Table 1. Correlations obtained with low-density (LD) or genotype-by-sequencing (GBS) data were compared to those achieved with the medium-density (MD) array (triangles versus dots). Reference inbreeding were estimated using the intermediate base population 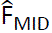.

When HBD levels were predicted for specific loci using low-density marker arrays, correlations with reference values decreased for all methods, more so when the reference base population was more distant. LocalNgsRelate and GERMLINE were two exceptions, as their correlations increased for several scenarios. Conversely, methods such as phasedibd, TRUFFLE or GIBDLD were more affected, with greater reductions in accuracy, whereas hap-IBD was relatively robust. Despite these variations, IBD_Haplo15c and ZooRoH performed best in all scenarios. Although hap-IBD was now the best rule-based method, it systematically achieved lower accuracy than the best model-based approaches.

For genome-wide predictions using GBS data, the accuracy increased with the number of fragments and their size, both of which affect the number of markers available. However, there were some exceptions. For example, the correlations were constant for ZooRoH-25, while they were identical for GBS-30K and GBS-50K when using IBD_Haplo15c. The multiple HBD class ZooRoH-125 performed better than these two approaches (ZooRoH-25 was better only when predicting recent HBD levels using GBS-15K), whereas the performance of the single HBD class ZooRoH model was severely compromised when using GBS data. The performance of the model-based approaches using GBS data was inferior to that using the MD array. Similarly, more accurate predictions were obtained using an LD array rather than a GBS-15K panel. On the contrary, rule-based methods such as phasedibd, hap-IBD and GERMLINE performed better with GBS-30K or GBS-50K panel than with MD arrays. These three methods are somewhat peculiar in that the highest accuracies were obtained using the LD and GBS-50K panels. Importantly, phasedibd was the most accurate method in almost all evaluated configurations (i.e. with the GBS marker densities and with different reference HBD levels), while hap-IBD was systematically better than model-based approaches with GBS-30K and GBS-50K panels. PLINK ROH had poor accuracies with GBS-15K, but its performance was good with GBS-50K. As a result, three rule-based methods achieved the highest genome-wide prediction accuracies when using the GBS-50K panel. Despite their lowest accuracies with GBS data, ZooRoH (multiple HBD class models) and IBD_Haplo15c remained among the best methods, still outperforming other model-based approaches such as LocalNgsRelate and GIBDLD. Interestingly, the ranking of the pedigree-based predictor increased, especially when fewer markers were available (GBS-15K or GBS-30K) or when predicting recent HBD levels. It even became the best or second best in some configurations.

When using GBS data for locus-specific predictions, accuracy increased systematically with the number of markers in the panel. It is important to note that the locus-specific accuracy was evaluated at the marker positions. For GBS data, this corresponds to small fragments with high marker density, and accuracy in less marker dense regions is not evaluated. Overall, ZooRoH models with multiple HBD classes achieved the highest accuracies, often followed by IBD_Haplo15c. Phasedibd or hap-IBD were regularly the next best methods, sometimes better than IBD_Haplo15c. PLINK-ROH performed poorly with GBS-15K or GBS-30K, but like phasedibd and hap-IBD, its ranking increased when more markers were available. At the locus-specific level, ZooRoH-1R was less affected by the use of GBS data. It was still significantly less efficient than the other ZooRoH models or IBD_Haplo15c when a young or intermediate reference base population was used, but it became the best approach for comparisons with the most distant base populations. Interestingly, ZooRoH-125 was best and ZooRoH-25 second best with the intermediate base population, whereas the opposite was observed with the recent base population.

When locus-specific prediction accuracy was assessed using AUC (Figure S9-10), ZooRoH with multiple HBD classes and IBD_Haplo15c were consistently the best across different scenarios. The only exception was the prediction of recent HBD levels using GBS-30K or GBS-50K panels, where hap-IBD resulted better than IBD_Haplo15c. Interestingly, for distant base populations, the highest AUC values were obtained with GBS-50K, whereas for recent HBD levels, higher values were obtained with the LD marker array.

## Accuracy of predictions when founder allele frequencies are used

Ideally, methods that use estimated AF in their model should work with founder AF, but these are rarely available. The DAMONA data set provides the opportunity to estimate these founder AF and to assess the impact of using founder AF versus sample AF in these methods.

For genome-wide prediction of HBD levels (Figure 6), the use of founder AF dramatically increased the correlations obtained with SNP-by-SNP approaches. Significant improvements were also observed for IBD_Haplo9c (using unphased data), which achieved similar accuracy to IBD_Haplo15c (using phased data). When using commercial genotyping arrays, the performance of the ZooRoH models, IBD_Haplo15c and LocalNgsRelate did not change with founder AF (we only observed slight increases with ZooRoH-1R). With GBS data, the multiple HBD classes ZooRoH model and LocaNgsRelate were not affected by the change in AF, whereas large and modest improvements were observed for ZooRoH-1R and IBD_Haplo15c, respectively. Nevertheless, ZooRoH-1R was still less accurate than the other ZooRoH models. For locus-specific predictions, the AF used had little effect, and we did not observe any changes in performance (Figure 6).

**Figure 6.**
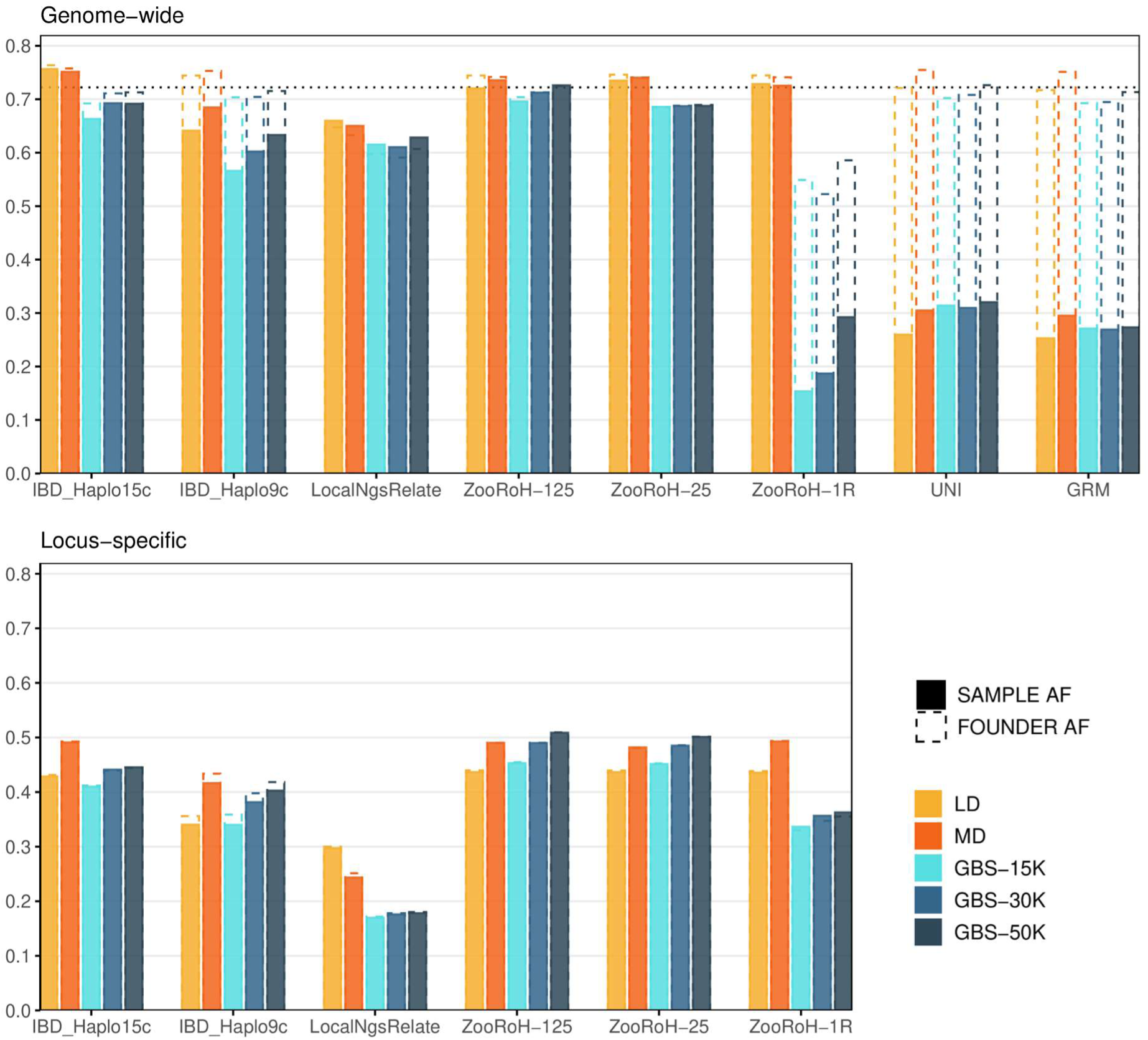
Impact of using founder versus sample allele frequencies on correlations between predicted and reference genome-wide and locus-specific HBD levels in the DAMONA cattle data set, for the methods described in Table 1 that accept external allele frequencies as input. Reference inbreeding were estimated using the intermediate base population 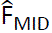. Correlations are shown for different marker panels including low density (LD), medium density (MD), and GBS panels with different number of markers. Correlations are compared for predictions computed using founder versus sample allele frequencies. The horizontal dashed line is the correlation obtained with the pedigree.

### Direct estimation of relatedness between the parents

The expected inbreeding coefficient of an individual is equal to the relatedness coefficient between its parents. Similarly, the prediction of HBD segments in the offspring is closely related to the identification of IBD segments between the parents. Thus, our evaluation procedure also provides indirect information on the accuracy of estimation of IBD segments and relatedness between parents. In simulated data, it is possible to directly assess the efficiency of the same methods for estimating relatedness. We therefore took advantage of our simulations to compare the accuracy of relatedness measures (see Figure S13). The ranking of the methods was the same as for the prediction of HBD levels, but with higher levels of accuracy and smaller differences between methods. The higher accuracy may be due to the fact that there is unpredictable random variation associated with Mendelian sampling in predicting HBD levels, a feature not present in kinship estimation. Interestingly, the pedigree-based approach was also less accurate for kinship because it can only estimate expected relatedness, whereas molecular-based estimators estimate realized relatedness.

## Discussion

### Evaluation with real and simulated data

We herein used simulations and empirical comparisons with two real data sets, corresponding to a typical livestock species and a wild population in a conservation program, to evaluate the accuracy of different approaches used to predict HBD levels in future offspring of a pair of genotyped parents, an important information for the management of these populations. The two types of data set we used are complementary. The true HBD levels are known in the simulations, whereas more realistic conditions are obtained with real data sets. These include levels of relatedness in the sample, past demographic history, genome and LD structure, distribution of AF, effects of past and ongoing selection, levels of genotyping and phasing errors, etc. Evaluations of prediction methods on both types of data showed the same trends in terms of ranking the methods, indicating that the empirical analysis on real data was informative. The two real data sets had different structures. MW had higher inbreeding levels and a smaller sample size, that might affect the phasing efficiency and the accuracy of estimation of AFs for example. These would be the typical field conditions in conservation genetics or studies of wild fauna. The use of small samples might also increase the role of random variation in method comparisons. The cattle population had the advantage to contain more individuals that were whole-genome sequenced, providing more information on the true inbreeding levels. In general, important information is available in livestock populations, including deep and accurate pedigrees, a good reference genome and genetic maps, that are important for methods identifying IBD segments (see also Bosse et al., 2015). Genotyping marker arrays are also available at different densities and the reference genome allowed us to simulate GBS data, although these were probably cleaner than real GBS data. This illustrates that data available from livestock populations might be useful to study techniques that will be applied in molecular ecology. Here, we had access to a relatively large sequenced pedigree including many trios, as needed for our study.

### Relative performance of evaluated methods

Using these data sets, we evaluated 16 methods for predicting HBD levels in the offspring of genotyped parents. Importantly, we showed that this evaluation is also informative about the accuracy of the methods for estimating relatedness and identifying IBD segments. Such estimators are useful in more applications than the prediction of HBD levels, including, for example, diversity management, selection, control for population structure, etc. In terms of accuracy, IBD_Haplo15c and multiple HBD class ZooRoH models (ZooRoH-MixKL) were consistently among the most efficient when evaluated across different scenarios, on simulated and real datasets. IBD_Haplo15c is the most complex model-based approach as it models the 15 identity states, while ZooRoH-MixKL, designed to estimate HBD levels within individuals, conceptually has two main states (HBD and non-HBD) but adds an additional layer of complexity by partitioning HBD into different length-based classes. IBD_Haplo15c was generally better at predicting recent HBD levels, whereas ZooRoH-MixKL was more efficient when the base population was more distant. Still, it was possible to shift ZooRoH towards predicting recent HBD levels by selecting the appropriate HBD classes, this approach being more efficient for locus-specific prediction. Both methods outperformed the best rule-based methods, especially when probabilities were useful (i.e. with ROC curves) and information was reduced, at lower marker density and for locus-specific predictions. This behavior has previously been described for reduced marker panels in Lavanchy and Goudet (2023), at lower marker densities (Solé et al. (2017); Druet et al. (2020)) and for locus-specific estimation of HBD levels (Alemu et al., 2021). Such locus-specific predictions could be useful for managing recessive deleterious alleles that cause genetic defects, loci that are major contributors to inbreeding depression, predicted harmful mutations, or for maintaining heterozygosity at specific loci that should have high diversity, such as the major histocompatibility complex. This could be particularly valuable in managing populations derived from a few founders with fully characterized genomes, including at deleterious loci. This information could also be used to give more weight to regions predicted to contribute to genetic load (Bertorelle et al., 2022), as neutral diversity may not always be representative of extinction risk (Teixeira and Huber, 2021). Locus-specific HBD predictions could also be used to manage inbreeding and diversity at all loci simultaneously, using algorithms that optimize all loci together. For genome-wide HBD predictions, phasedibd, PLINK-ROH and, to a lesser extent, hap-IBD, were also very accurate. Nevertheless, the performance of PLINK-ROH and hap-IBD decreased in some configurations, for example with LD and GBS-15K panels for PLINK-ROH which may be even less accurate in the presence of genotyping errors. Phasedibd performed well in all scenarios and thus represents an excellent option for genome-wide predictions, combining accuracy and computational efficiency. Here we have focused on small samples of interest in conservation genetics and wildlife, where model-based approaches are still applicable, but this may not be the case for larger data sets or at higher marker densities (biobanks and large livestock populations). However, these rule-based approaches were less efficient for locus-specific predictions, especially with the genotyping arrays, while the efficiency improved at higher marker densities such as with the GBS-50K panel. The other methods evaluated were less accurate in most scenarios, although they could perform well in some rare exceptions. In addition, many of them had high variability and therefore performed particularly poorly in some configurations. The SNP-by-SNP approaches did not perform well in all configurations, especially with sample AF, and only provide genome-wide predictions, but they have the advantage of being applicable with sub-optimal genome assemblies or without a genetic map.

### Method features affecting predictions accuracy

From these comparisons we learned several properties of the prediction methods. First, the use of phased data achieved higher accuracy than the use of genotypes, despite possible errors introduced during the phasing process, consistent with the findings of Gómez-Romano et al. (2016). This was also the case for (real) small data sets, where phasing errors are expected to be more common. Interestingly, we observed no differences in accuracy using true versus estimated haplotypes at the genome-wide level, and only minor differences for locus-specific predictions (Figure S14). However, accurate phasing is not always possible, e.g. when physical or genetic marker maps are not available or with low-fold sequencing data (genotypes are not unambiguously known). Among the methods evaluated, LocalNgsRelate was the only one to handle low-fold sequencing data. This feature could also be added to other model-based methods using unphased data (e.g. IBD_Haplo9c). Second, several methods, including the SNP-by-SNP approaches, were sensitive to the AF used. In agreement with theoretical expectations and the study by Caballero et al. (2022), better estimators were obtained with founder AF. Interestingly, the performance of IBD_Haplo9c (using unphased data) increased close to that of IBHaplo15c (using phased data). This suggests that when phasing accuracy is compromised, such as with smaller datasets or low-fold sequencing data, methods that work with unphased data may be a good option if founder AF are available, but this is unfortunately rarely the case. Here we used the gene content approach, which requires a relatively large genotyped sample and a deep pedigree, which are rarely available in conservation genetics or wildlife. Conversely, it also shows that when founder AFs are not known, it is better to rely on a method using phased data. Overall, several methods were robust to the AF used, including obviously the rule-based approaches, but also IBD_Haplo15c and ZooRoH-MixKL. It should be noted that the AF used had only a marginal effect on the accuracy of locus-specific HBD predictions, which depend more on the homozygosity of the markers around the target position. Thirdly, the 3-STATES models were not among the best methods. This suggests that the assumption that the parents were non-inbred is not optimal in populations with small N_e_. Locus-specific coancestry and predicted HBD levels are indeed reduced if we ignore that the haplotypes from one or both parents are IBD (e.g. the maximum value drops from 1 to 0.5), and this can have a significant impact in inbred populations. Fourth, we also learned that the use of fixed parameters defining the frequency of IBD (as in IBD_Haplo) or the length of IBD segments (as in IBD_Haplo or ZooRoH-MixKL) did not result in worse performance than the estimation of these parameters (as in GIBDLD or LocalNgsRelate). Finally, we observed that multiple HBD classes ZooRoH models were more robust to marker density or used AF than a ZooRoH-1R model. This is in agreement with the observation that pruning strategies are recommended with single HBD-class HMM (Leutenegger et al., 2003; Narasimhan et al., 2016; Vieira et al. 2016), whereas this is not necessary with a ZooRoH-MixKL model, as we have previously shown (Druet and Gautier, 2017). Nevertheless, the ZooRoH-1R performed well with the LD and MD genotyping arrays.

### Performance of pedigree-based predictions

We observed that in some configurations, pedigree-based predictions of HBD levels performed well compared to methods using molecular data. This is different from estimating individual levels of inbreeding, where genomic estimators proved superior (Keller et al., 2011; Wang, 2016). In this situation, molecular data allow estimation of realized HBD levels, whereas the pedigree-based approach is limited to expected levels because it can not predict Mendelian sampling. In the prediction context, both approaches are unable to predict Mendelian sampling and therefore achieve more similar accuracies. The pedigree-based approach will be more competitive for predicting recent HBD levels (since pedigrees only capture recent generations), and when the accuracy of molecular-based approaches decreases, such as with LD marker arrays, some GBS panels, unavailable founder AF (for some methods) and inaccurate phasing. These results are in agreement with Woolliams et al. (2022), who suggested that the pedigree-based approaches could be a good option for genetic diversity management. However, the pedigree-based estimators require that the genealogy is accurately and deeply recorded, which is not always possible in wildlife populations for example. This argues for recording pedigrees as well as possible for population management, even when molecular data are available (Galla et al., 2021).

## Acknowledgements

Tom Druet is Research Director from the Fonds de la Recherche Scientifque – FNRS (F.R.S-FNRS). Computation were carried out using the supercomputing facilities of the ‘‘*Consortium d’Equipements en Calcul Intensif en Fédération Wallonie-Bruxelles*’’ (CECI), funded by the F.R.S-FNRS.

