## Supplementary Figures for "Genomic prediction of individual inbreeding levels for the management of genetic diversity in populations with small effective size"

**Table S1.** Arguments used to run the methods

| Software | Arguments |
|---|---|
| IBD_Haplo | With haplotypes: those in gold standard files phased_2009.par and phased_2011.par, with the original and modified transition matrix, respectively<br>With genotypes: those in gold standard files unphased_2011.par and unphased_2009.par |
| IBDLD | Default options, except -ibd 90 --ibdtxt |
| LocalNgsRelate | Default options, except l = 0 |
| RZooRoH | For the Mix7R: zoomodel(K=7, base=5, layers=TRUE)<br>For the 1R : zoomodel(K=2, predefined=FALSE, layers=FALSE) |
| phasedibd | Default options, except L_m=50 and L_f=2 |
| hap-IBD | Default options |
| GERMLINE | Default options, except -min_m 2 -haploid |
| TRUFFLE | Default options, except --segments --maf -1 --missing 1 --mindist -1 --L 2 --ibs1markers 100 --ibs2markers 50 |
| PLINK | Default options, except --homozyg-kb 2000 --homozyg-snp 40 --homozyg-density 100 --homozyg-gap 500 --homozyg-het 0 --homozyg-window-snp 40 --homozyg-window-het 0 |
| Refined IBD | Default options |
| GCTA | Default options, except maf 0 and make-grm-alg 0 (for UNI) or make-grm-alg 1 (for GRM) |

**Table S2.** Correlation (mean and confidence interval across 100 replicates) between the true genome-wide inbreeding levels and the genome-wide inbreeding estimated from 25,000 SNP genotypes with either RZooRoH (running a Mix7R model with base rate of 5 and defining *F* as the HBD proportion accumulated up to class with R=125) or PLINK (as the proportion of the genome in ROH of at least 2Mb). True inbreeding levels were defined with respect to a recent (YOUNG), intermediate (MID) and more distant (TOT) base population.

| Scenario | *F* estimator | Reference inbreeding levels | | |
|---|---|---|---|---|
| | | TOT | MID | YOUNG |
| MODF | HBD | 0.9953 ± 0.0004 | 0.9937 ± 0.0006 | 0.9625 ± 0.0035 |
| | ROH | 0.9818 ± 0.0017 | 0.9852 ± 0.0014 | 0.9710 ± 0.0026 |
| HIGHF | HBD | 0.9959 ± 0.0006 | 0.9959 ± 0.0006 | 0.9891 ± 0.0015 |
| | ROH | 0.9771 ± 0.0043 | 0.9778 ± 0.0042 | 0.9736 ± 0.0048 |

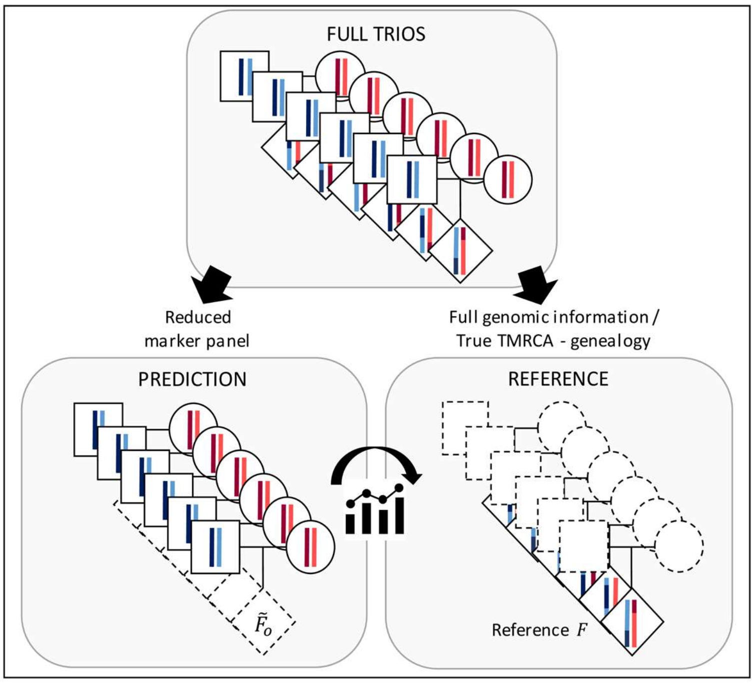


Figure S1. Validation design to evaluate the accuracy of predicting levels of homozygosity-by-descent (HBD) in future offspring based on parental genotypes in populations with small $N_e$. Our validation design relied on the use of sequenced or genotyped trios (sire-dam-offspring). Six trios including a sire (square), a dam (circle) and an offspring (diamond) are represented. The two colored vertical lines in either the sire (in blue) or the dam (in red) represent their two haplotypes. In this design, we divided the data in two: parents for prediction and offspring to estimate reference levels. In the prediction set, we masked the genotypes of the offspring (dashed lines) and used only the parental genotypes to perform the predictions of $\tilde{F}_o$ with the 16 evaluated methods.

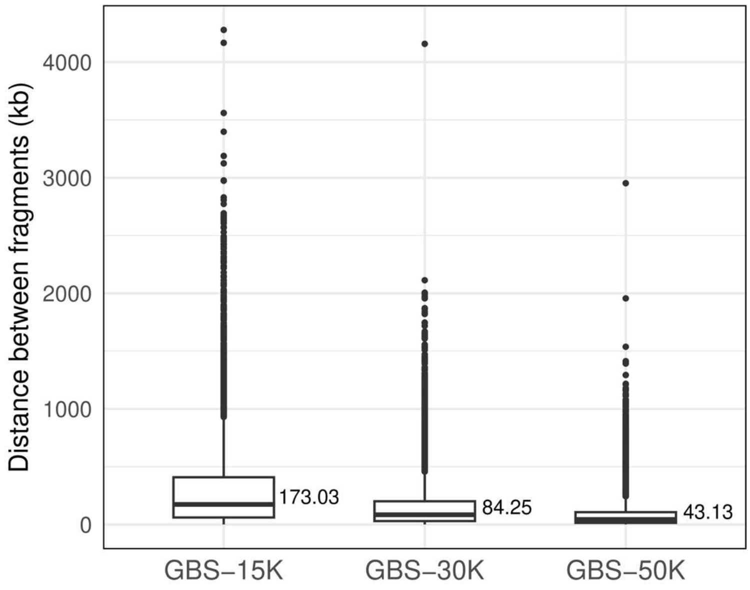


Figure S2. Distance (in kb) between consecutive fragments in the genotyping-by-sequencing (GBS) data with 56,098 SNPs (GBS-50K), 31,339 SNPs (GBS-30K) and 15,493 SNPs (GBS-15K). The GBS data was obtained by *in silico* digestion of the bovine reference genome and using the DAMONA cattle data set.

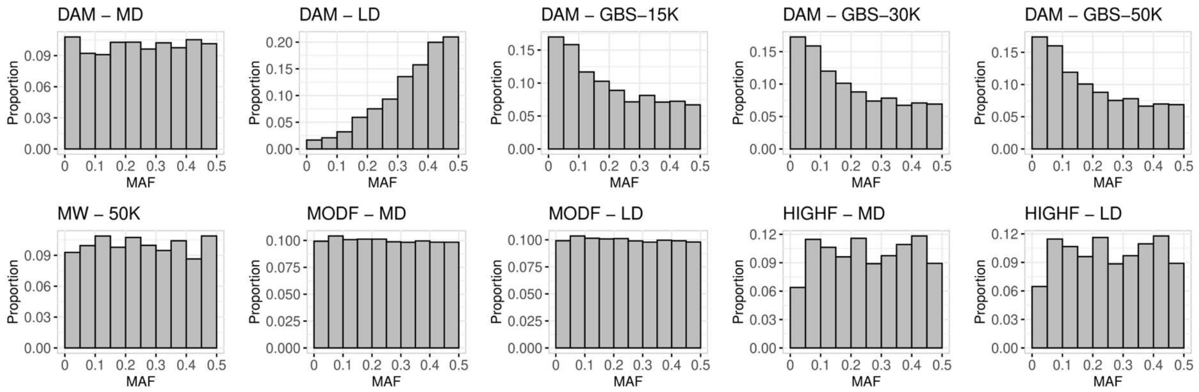


Figure S3. The distribution of minor allele frequency (MAF) observed in the four data sets for the different marker panels. Top panels show the distributions for the DAMONA cattle data set with either a medium density (DAM - MD), a low density (DAM - LD) or genotyping-by-sequencing panel with different number of markers (DAM – GBS-15K, DAM – GBS-30K, DAM – GBS-50K). Bottom panels show the distributions for the Mexican Wolf data set with a medium density array (MW - 50K), and for the moderately inbred (MODF) and highly inbred (HIGHF) simulated scenarios, with either a medium density (MODF - MD and HIGHF - MD) or a low density (MODF - LD and HIGH - LD) panel.

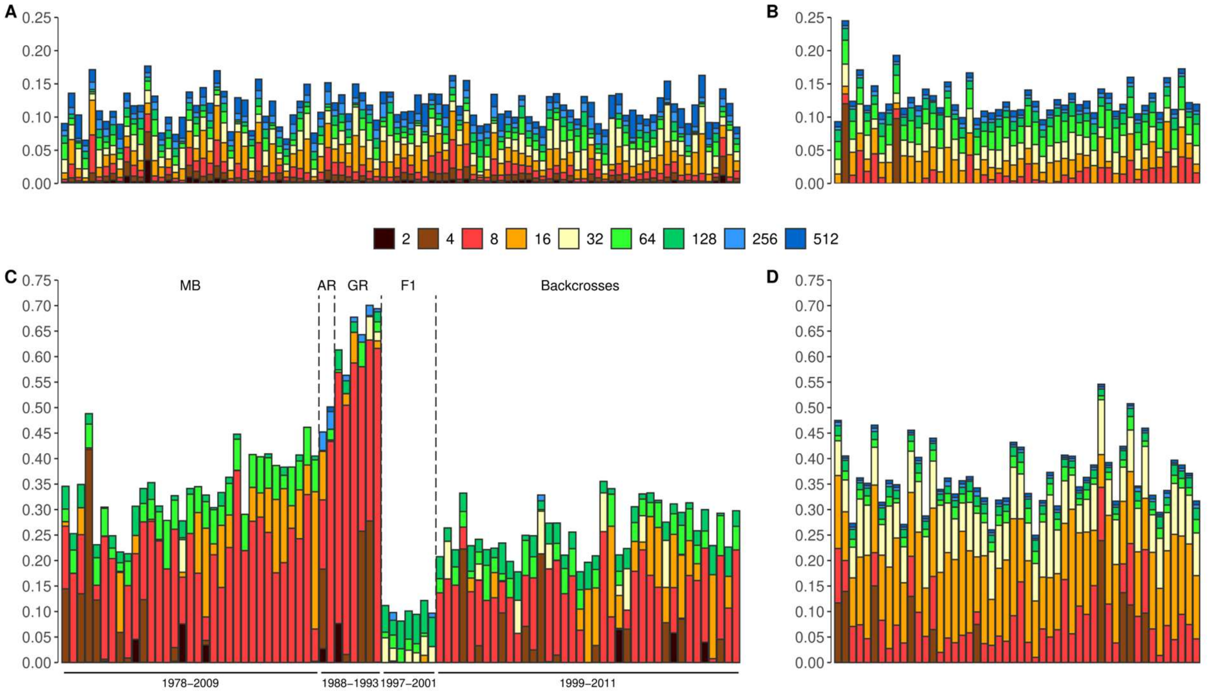


Figure S4. Autozygosity levels and age-based partitioning of HBD for the different data sets used in our study. A: DAMONA cattle data set, B: moderately inbred (MODF) simulated population, C: Mexican Wolf data set, and D: highly inbred (HIGHF) simulated population. Mexican wolves are sorted chronologically within each captive lineage (McBride (MB), Aragón (AR) and Ghost Ranch (GR)), crosses (F1) and backcrosses. For the real data, this was done by running RZooRoH with a model of 14 HBD classes with rates $R_c = \{2,4,8,\ldots,16384\}$ at the highest marker density available. These classes were chosen to obtain a finer classification of the partitioning. For the simulated data, the partitioning was obtained from the true TMRCA of the HBD segments, with HBD classes similar to those used for the real data. The partitioning was presented for HBD classes with a rate $R_c \leq 512$ (approximately 250 generations), as shorter HBD segments can not be reliably identified with the marker density available for the MW. We observed similar levels of inbreeding and partitioning of HBD in the real data set and the corresponding simulations. The average levels of inbreeding were equal to 0.12 in the cattle data and 0.13 in the MODF scenario, while these values were equal to 0.31 in the MW and to 0.36 in the HIGHF scenario. In the MW population and the HIGHF scenario, the recent HBD classes with rates of 4, 8 and 16 (corresponding to ancestors present 2 to 8 generations ago) had larger contributions to the total HBD. Nonetheless, a few individuals in the cattle or MODF populations showed also high levels of recent inbreeding. In the MW, the HBD partitioning reflected the known history well. In the three different lineages, we observed extremely high levels of inbreeding (on average 0.34 in MB, 0.47 in AT and 0.65 in GR), with lower levels estimated in the MB lineage derived from more ancestors (3 instead of 2). As expected, the level of inbreeding within these closed populations increased with birth year, but autozygosity shifted to more ancient HBD classes. This makes sense as these more recent individuals are more distant from the founders of the lineages. The first generation of crossbred individuals had almost no recent inbreeding, confirming that three lineages are genetically distant. The level of inbreeding tended to increase in subsequent generations of crossbreeding, and we again observed a shift towards more ancient HBD classes for more recently born individuals. Overall, we observed moderate to high levels of inbreeding in both real data sets, including HBD segments associated with recent ancestors. The simulated data closely matched the levels and partitioning observed in the corresponding real data. Comparisons also illustrate that dairy cattle and Mexican wolf populations, and the corresponding MODF and HIGH simulations, provide complementary scenarios.

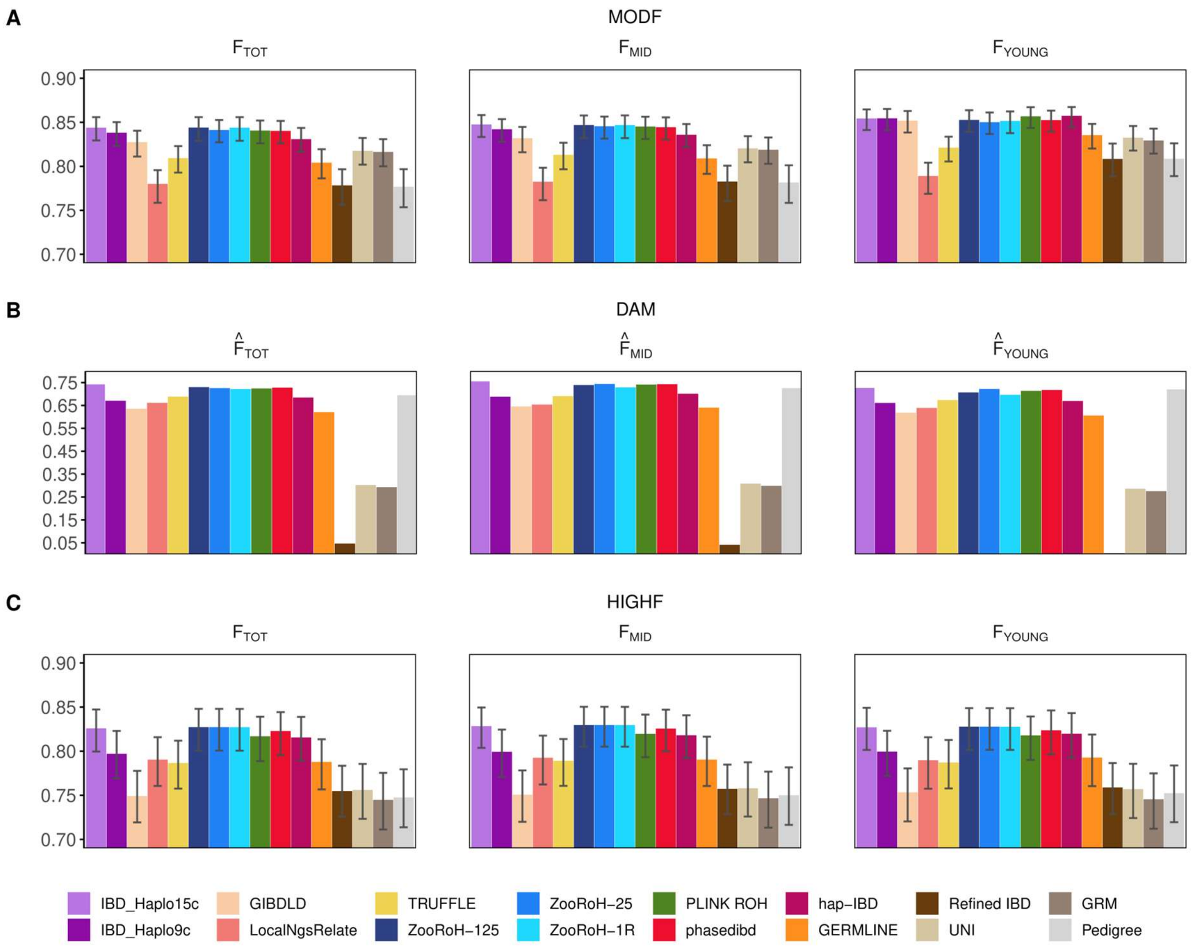


Figure S5. Correlations between predicted and reference genome-wide levels of HBD for the 16 methods compared in three scenarios, using a medium density array. Methods and their abbreviation are described in Table 1. A) moderately inbred simulated population (MODF); B) DAMONA cattle data set (DAM); C) highly inbred simulated population (HIGHF). The reference levels were defined for either a recent, an intermediate or a distant base population (approximately 15, 50 or 500 generations ago, respectively). Mean and 99% confidence intervals are shown for the simulated scenarios.

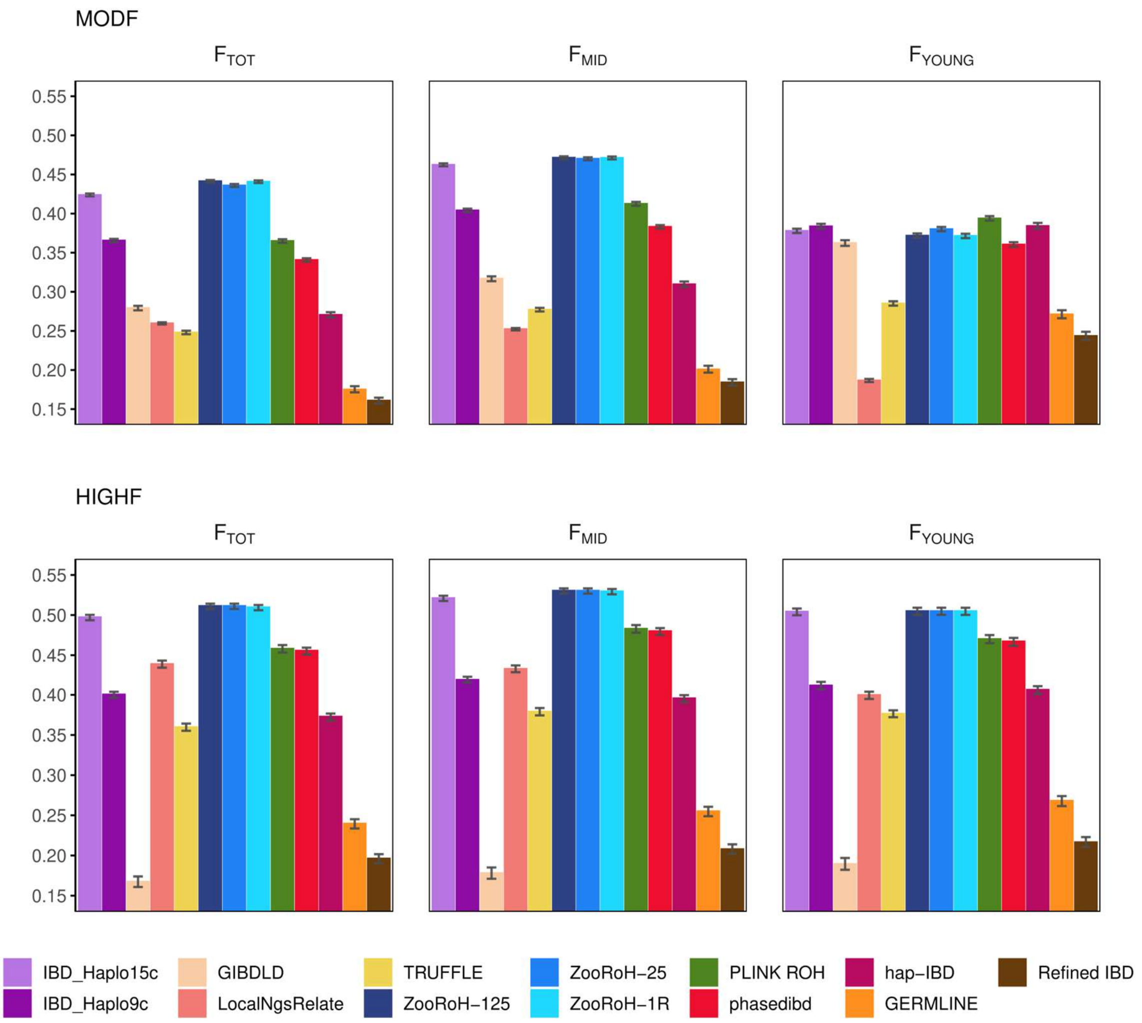


Figure S6. Locus-specific accuracy of 13 HBD prediction methods in the simulated data sets using a medium density array. Methods and their abbreviation are described in Table 1. Accuracy was assessed using correlations (mean and 99% confidence intervals) between predicted and reference locus-specific HBD levels in the MODF and HIGHF scenario. The reference levels are the true HBD status at every marker position and defined using either a recent (15 generations ago - $F_{YOUNG}$), an intermediate (50 generations - $F_{MID}$) or a more distant (500 generations ago - $F_{TOT}$) base population.

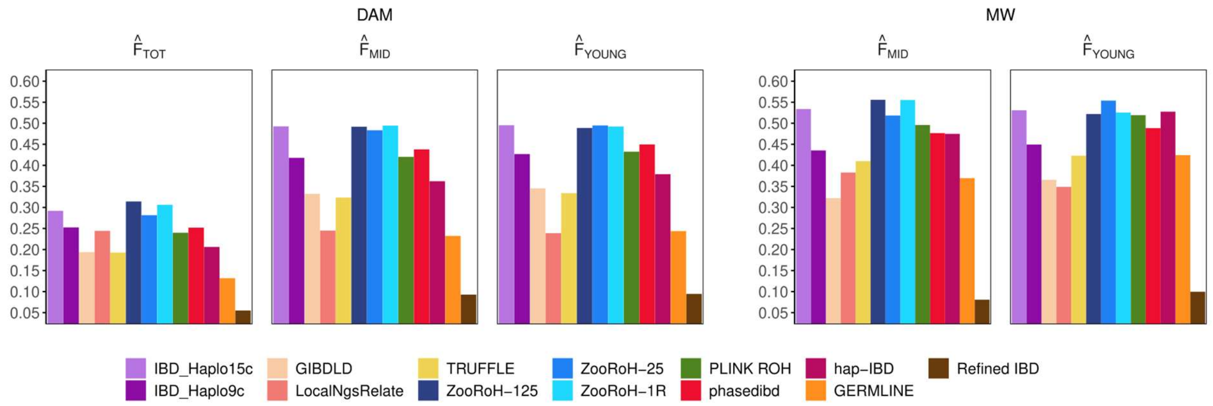


Figure S7: Correlations between predicted and reference locus-specific HBD levels in the DAMONA cattle (DAM) and the Mexican Wolf (MW) data sets. Reference inbreeding were estimated using three different base populations (recent - $\hat{F}_{YOUNG}$, intermediate - $\hat{F}_{MID}$ and distant - $\hat{F}_{TOT}$).

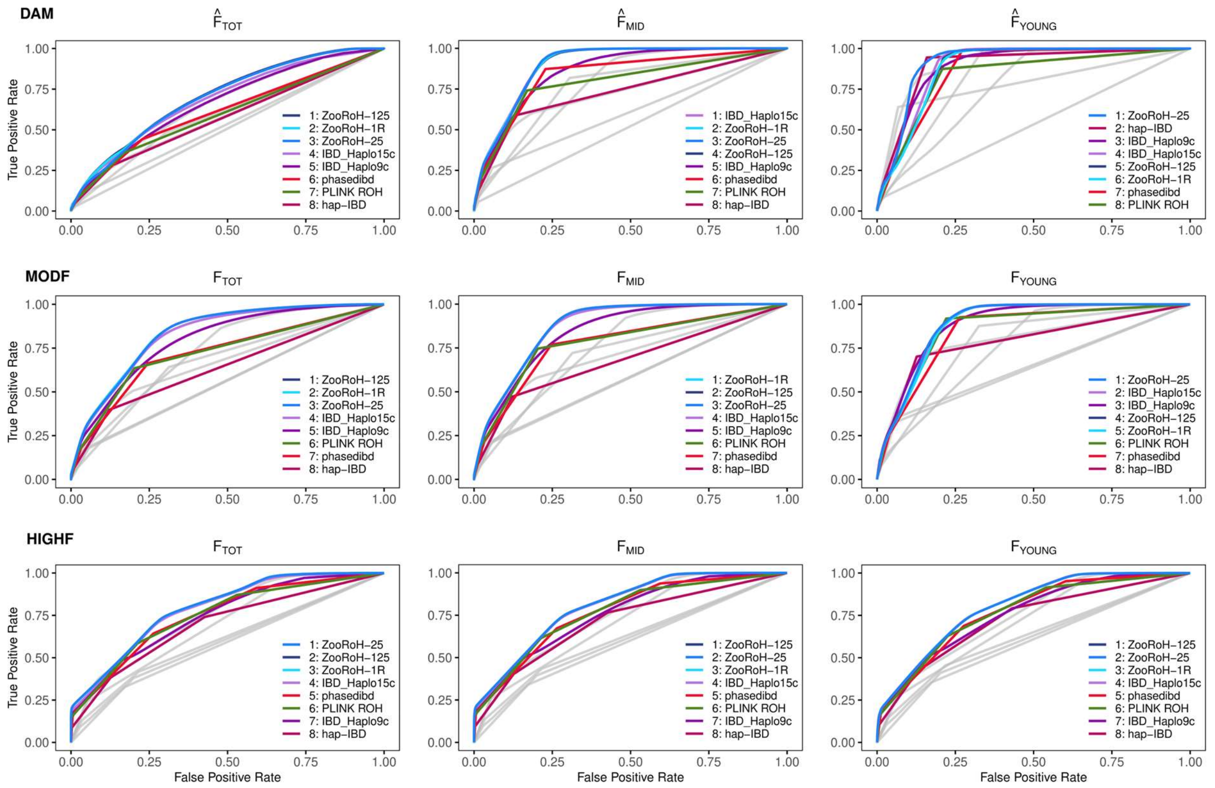


Figure S8. Receiver operating characteristic (ROC) curves obtained for 13 methods with a medium density panel for the DAMONA cattle data set (DAM) the moderately inbred (MODF) and the highly inbred (HIGHF) simulated scenarios. Reference inbreeding were estimated using three different base populations set approximately at 15 ($F_{YOUNG}$), 50 ($F_{MID}$) and more than 500 ($F_{TOT}$) generations in the past. Curves obtained with ZooRoH, IBD_Haplo, phasedibd, hap-IBD and PLINK ROH are highlighted and their relative ranking in terms of AUC is also specified.

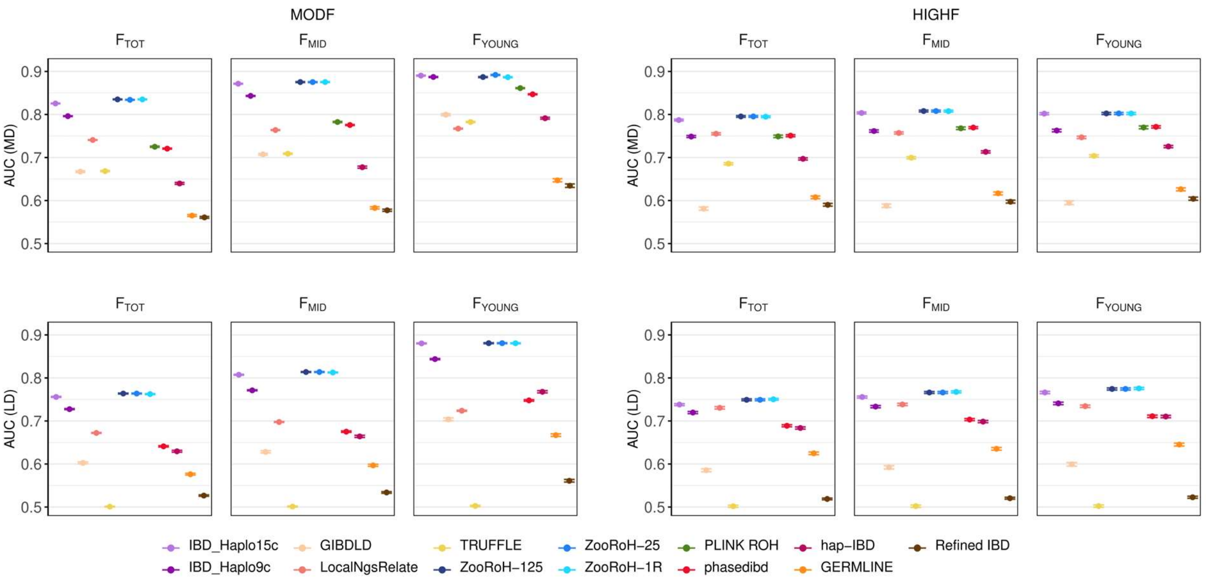


Figure S9. The area under the ROC curve (AUC) obtained for 13 methods evaluated for locus-specific performance in the moderately inbred (MODF) and highly inbred (HIGHF) simulated scenarios, using a medium density (MD) or a low density (LD) panel. Results include mean correlation and 99% CI across 100 replicates. The true HBD status was the reference inbreeding, defined using either a young (15 generations - $F_{YOUNG}$), an intermediate (50 generations - $F_{MID}$), or a distant (500 generations - $F_{TOT}$) base population.

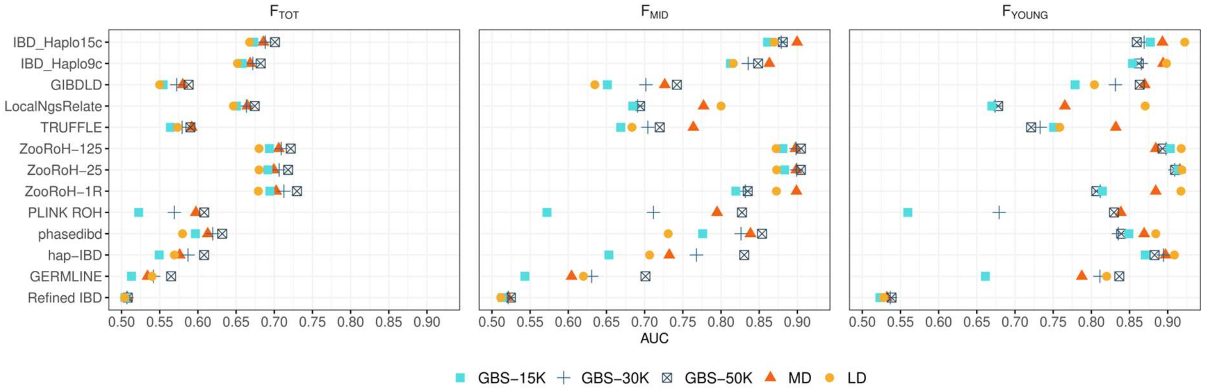


Figure S10. The area under the ROC curve (AUC) obtained for 13 methods evaluated for locus-specific HBD prediction methods using the DAMONA cattle data set and different marker panels: low density (LD), medium density (MD) and genotyping-by-sequencing (GBS) data with different number of markers. Reference inbreeding were estimated using three different base populations (recent - $\hat{F}_{YOUNG}$, intermediate - $\hat{F}_{MID}$ and distant - $\hat{F}_{TOT}$).

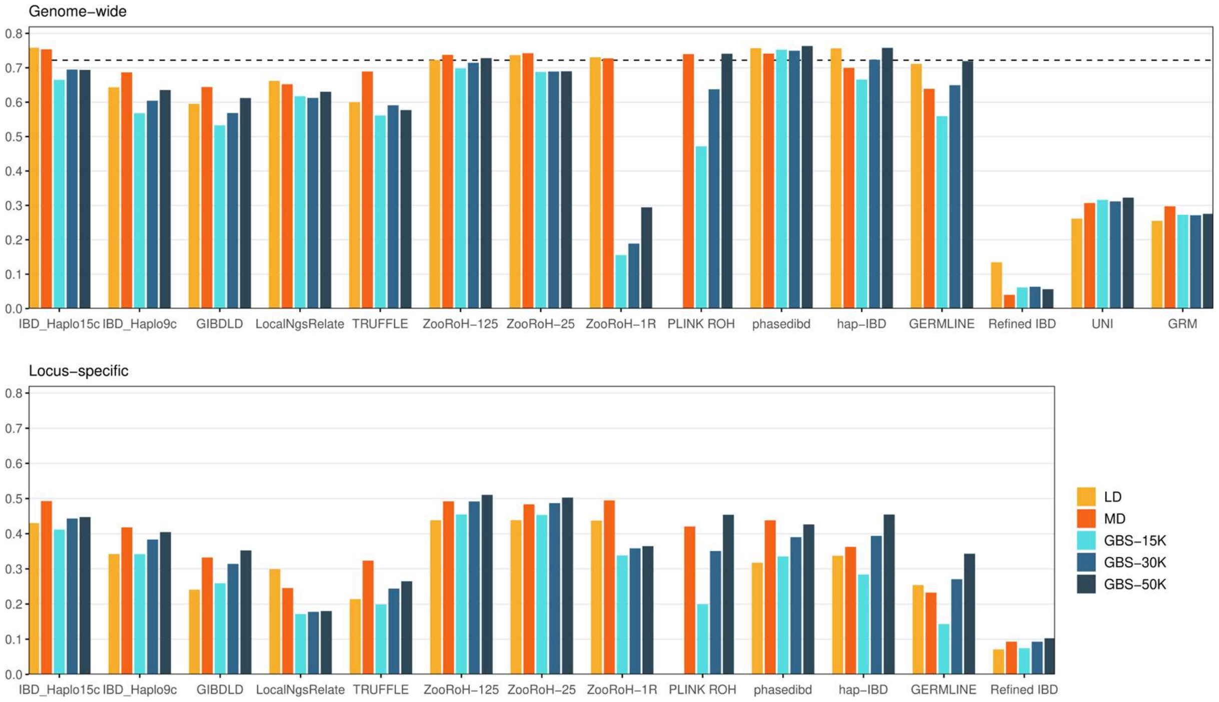


Figure S11. Correlations between predicted and reference genome-wide or locus-specific HBD levels for the 16 evaluated methods using the DAMONA cattle data set and reduced genotyping arrays including low density (LD), medium density (MD, red), and genotyping-by-sequencing (GBS) panels with different number of markers. Methods and their abbreviation are described in Table 1. Reference inbreeding were estimated using the intermediate base population $\hat{F}_{MID}$.

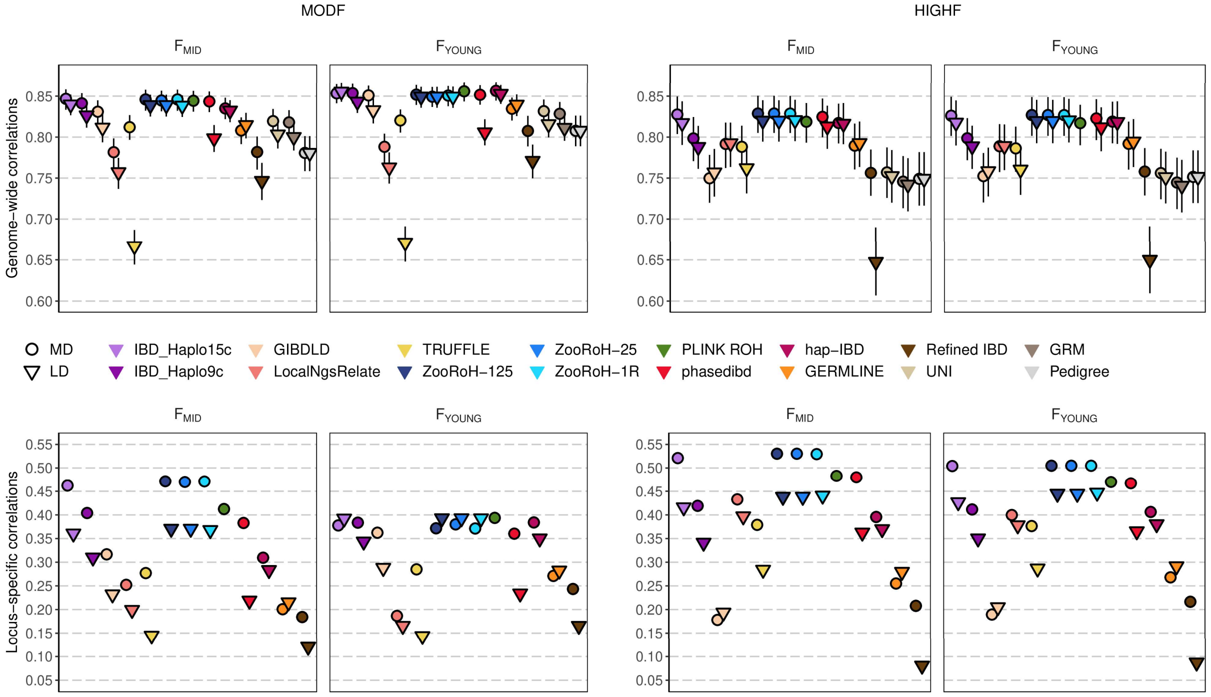


Figure S12. Correlations between predicted and reference genome-wide or locus-specific HBD levels for the 16 evaluated methods in the moderately inbred (MODF) and the highly inbred (HIGHF) simulated scenarios. Results, including mean and 99% confidence intervals, are shown for the predictions obtained with a low density panel (LD) and compared side-by-side against those obtained with a medium density panel (MD). The reference inbreeding was the true HBD proportion of the genome (or the HBD status at the locus), defined using either a young (15 generations - $F_{YOUNG}$), an intermediate (50 generations - $F_{MID}$), or a distant (500 generations - $F_{TOT}$) base population.

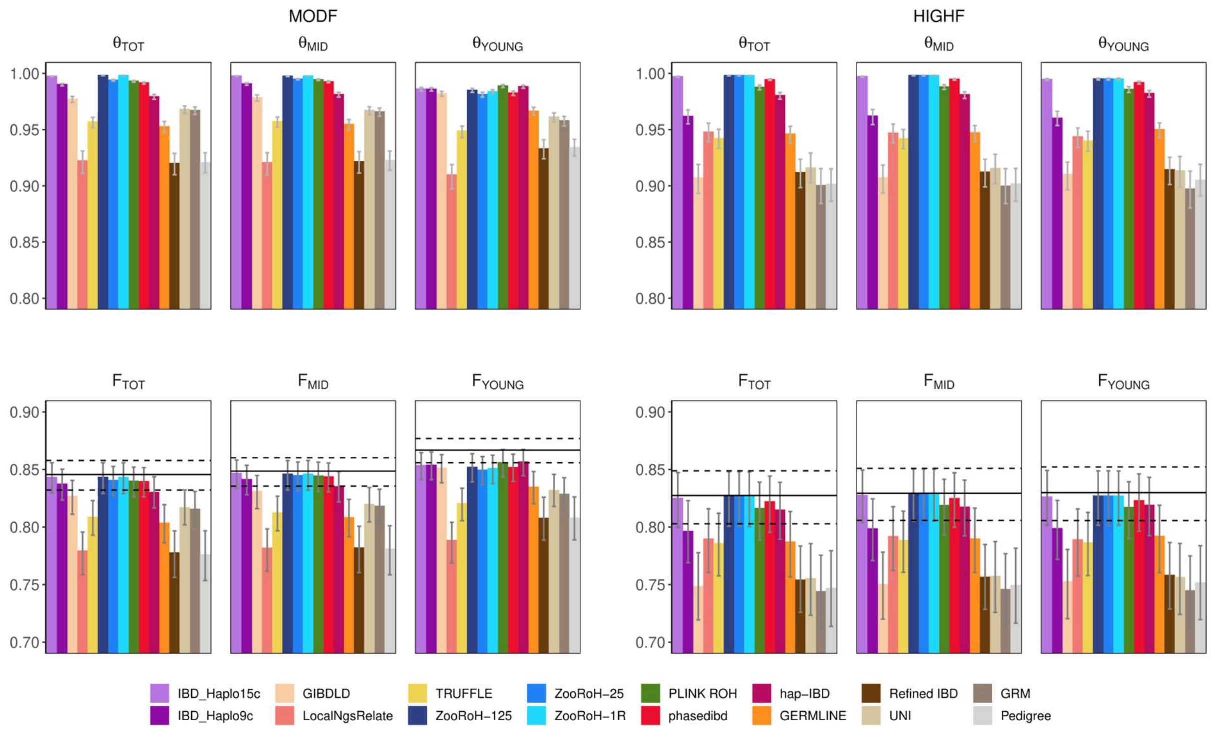


Figure S13. Correlations between estimated and true coancestry levels among pairs of parents ($\theta$). Coancestry levels were estimated as described in Figure 1 and using the 16 methods described in Table 1. Results, including mean and 99% confidence intervals, were obtained using the moderately (MODF) and highly inbred (HIGHF) simulation scenarios. The true coancestry levels were obtained using either a young (15 generations - $\theta_{YOUNG}$), an intermediate (50 generations - $\theta_{MID}$), or a distant (500 generations - $\theta_{TOT}$) base population. Correlations between predicted and true genome-wide levels of HBD in the offspring ($F_O$) are also provided to facilitate comparisons (bottom). The horizontal line represents the average correlations between the true coancestry between the parents and the true HBD level in the offspring, and indicates thus the maximum expected levels for HBD prediction (dashed horizontal lines represents confidence intervals).

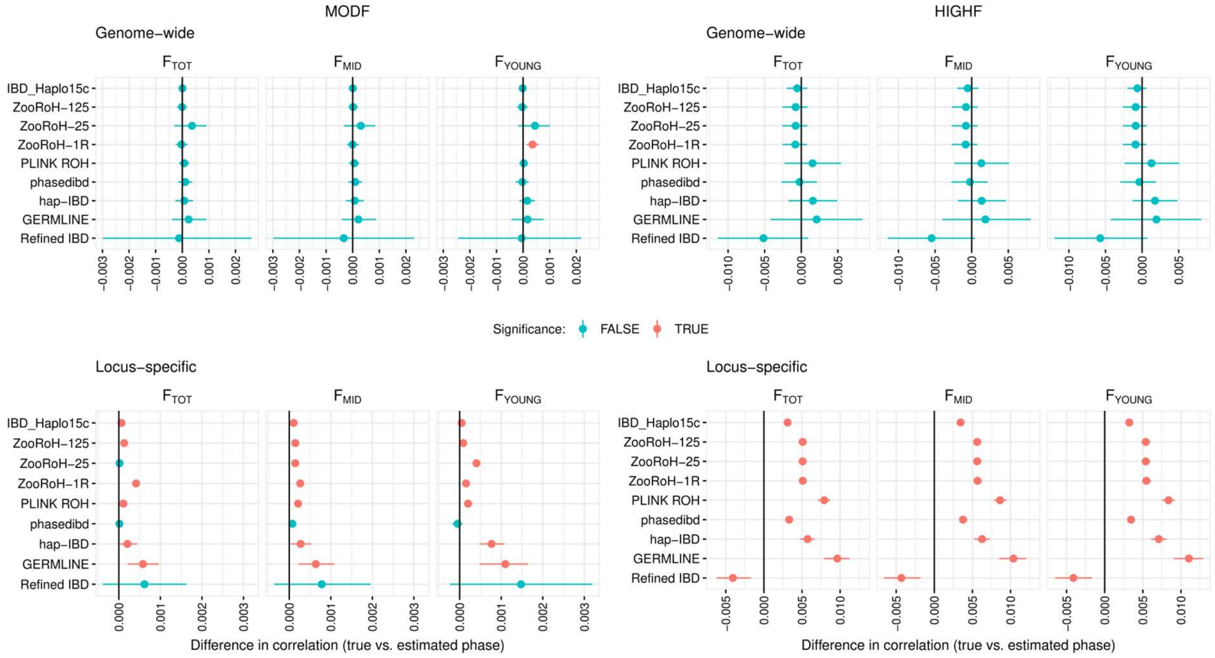


Figure S14. Impact of phasing accuracy on correlations between predicted and reference genome-wide (top) and locus-specific (bottom) levels of homozygosity-by-descent, in the moderately (MODF) and highly (HIGHF) inbred simulated datasets. Results are shown for prediction methods that use haplotypes. The values represent the mean and confidence interval across 100 replicates of the difference between the correlation computed using either the true haplotypes or the estimated haplotypes. The reference HBD levels are defined using either a young (15 generations - $F_{YOUNG}$), an intermediate (50 generations - $F_{MID}$), or a distant (500 generations - $F_{TOT}$) base population.
